# Transferrin receptor-mediated transport at the blood-brain barrier is elevated during early development but maintained across adult aging

**DOI:** 10.1101/2024.11.12.623253

**Authors:** Vanessa O. Torres, Michelle E. Pizzo, Darren Chan, Jason C. Dugas, David Huynh, David Joy, Eric K. Liang, Lily Sarrafha, Isabel Becerra, Roni Chau, Kylie S. Chew, Johann Chow, Timothy K. Earr, Nathalie Khoury, Kendra J. Lechtenberg, Amy W. Leung, Hoang N. Nguyen, Emmanuel S. Ojo, Elysia Roche, Hilda Solanoy, Mabel Tong, Raymond K. Tong, Kirk Henne, Joseph W. Lewcock, Ryan J. Watts, Meredith E. Calvert, Robert G. Thorne, Y. Joy Yu Zuchero

**Author notes:** **Corresponding authors:** Robert G. Thorne, PhD and Y. Joy Yu Zuchero, PhD, Biology Discovery, Denali Therapeutics Inc., 161 Oyster Point Blvd, South San Francisco, California 94080, and. Designates co-corresponding authors (Robert G. Thorne, Y. Joy Yu Zuchero).

## Abstract

Transferrin receptor (TfR)-mediated transcytosis across the blood-brain barrier (BBB) is a promising strategy to improve delivery of biologics to the central nervous system (CNS). However, it remains unclear whether age and aging-related diseases impact TfR expression and/or BBB transport capacity. Here, we used the TfR-targeted antibody transport vehicle (ATV^TfR^) to enhance CNS delivery in healthy mice and in the 5xFAD mouse model of Alzheimer’s disease (AD). Healthy neonates exhibited the highest vascular TfR expression and ATV^TfR^ brain exposure, whereas BBB transport capacity remained stable across adulthood. Additionally, neither TfR expression nor ATV^TfR^ brain uptake changed significantly in 5xFAD mice. Further, vascular TfR expression in AD patient brains was similar to age-matched controls, suggesting that TfR transport may be conserved for AD in humans. The elevated TfR-mediated brain delivery observed in early mouse development suggests the potential of added efficacy in utilizing TfR platforms for the treatment of early childhood diseases. Preservation of ATV^TfR^ transport in adult mice across healthy aging and in an AD model supports continued application of TfR platforms in age-related diseases.

## Introduction

The highly restrictive nature of the blood-brain barrier (BBB) poses a significant challenge for the delivery of many small molecules and nearly all large molecules into the central nervous system (CNS) (*1-3*). With only 0.01-0.1% of systemically administered IgG typically entering the CNS (*4*), the development of new IgG-based neurotherapeutics that utilize active transport mechanisms and receptor-mediated transcytosis (RMT) from the luminal (blood) to abluminal (brain) side of brain endothelial cells (BECs) has become a major research area (*4-6*). In particular, engineered binding to transferrin receptor 1 (TfR) has been demonstrated by multiple groups to significantly improve the efficiency of CNS large molecule delivery in rodents (*7-14*) and non-human primates (*14, 15*). Despite these promising efforts, it remains unclear whether and how healthy aging across a broad age range, as well as the presence of a neurodegenerative disease, e.g. Alzheimer’s disease (AD), might impact TfR-mediated transport at the BBB. During healthy adult aging, the BBB undergoes various structural, metabolic, inflammatory, and trafficking-related changes (*16-18*), in addition to the restructuring of the neurovascular unit (*19*). These changes may potentially alter TfR cycling rates and/or the endocytic machinery used to transport TfR across the BBB. Additionally, transcriptional and proteomic changes in BECs are well documented in the context of AD (*20-24*), which may further impact the delivery of TfR-targeted therapeutics to the CNS. Further, a potential increase in CNS barrier permeability due to compromised integrity and/or dysfunction of the brain barriers has been suggested in both healthy aging (*17, 19, 25*) and AD (*19, 26*). All of these factors have the potential to impact the effectiveness of RMT-based delivery of CNS drugs. Therefore, understanding how age and AD impacts TfR-mediated transport as well as CNS permeability in these settings will be critical to assessing the practical utility of TfR-based BBB transport platforms, a number of which are now under clinical evaluation (*27, 28*).

To date, only a few studies have evaluated TfR expression and TfR-mediated transport of IgG at the BBB in healthy aging and disease mouse models (*17, 29-31*). These studies have often produced varying observations, likely due to differences in methodology, mouse ages, genotype, TfR antibody designs (*17, 30*), and timepoints (*17, 30, 31*). The extent to which aging may impact parenchymal brain uptake of molecules using a TfR-based platform has remained unclear. Although transport of TfR ligands (i.e., iron and transferrin) across the BBB has been suggested to peak during early development (*32, 33*) and then decrease with adult aging (*32, 33*), the available data has been limited and often lacking in quantitative comparisons using validated methodology spanning early and late development, aging, and disease.

A TfR-binding antibody transport vehicle (ATV^TfR^), where the human IgG (huIgG) Fc is engineered to bind TfR and enhance delivery of therapeutics to the CNS has been previously reported (*14, 34*). Here, we utilized this well characterized ATV^TfR^ architecture (*14*) to investigate how healthy aging across a broad age range and disease progression in a mouse model of AD impacts brain TfR expression and BBB transport of endogenous IgG, standard antibodies, and ATV^TfR^. Multiple approaches were utilized to evaluate TfR expression and ATV^TfR^ transport in wild-type and TfR^mu/hu^ knock-in (KI) mice as well as in 5xFAD (familial AD) transgenic mice. TfR expression was also evaluated in post-mortem human brain tissue of AD and non-AD subjects to better understand the potential clinical translatability of the findings. Our results reveal that vascular TfR expression is significantly elevated in neonates when compared to adults and that ATV^TfR^ maintains the ability to significantly increase brain exposure of huIgG across a broad age range and in the presence of plaque pathology. The results also demonstrate limited brain uptake of endogenous IgG in both aged and 5xFAD mice, confirming BBB permeability to IgG is not significantly altered with aging and in this disease model. Taken together, our results demonstrate the maintenance of robust ATV^TfR^-mediated CNS delivery in both healthy aging and in 5xFAD mice.

## Results

### Total BEC TfR expression is highest in young mice and the accessible pool of vascular TfR remains consistent throughout adulthood

To determine whether TfR expression differs across a broad age range, we first evaluated total TfR expression in the brains of TfR^mu/hu^ KI mice ranging from postnatal day 2 (0.03 months; neonates) through 24 months of age. Evaluation of TfR protein expression in bulk brain lysate from the anterior hemisphere revealed that TfR expression is approximately 2-fold higher in neonates but remains consistent and stable through adulthood (**Fig. S1a, Fig. 1a**). Since the utility of TfR-mediated CNS drug delivery platforms rely on expression of TfR on the brain vasculature, we next asked whether age specifically impacts vascular TfR expression. Vascular segmentation (see Material and Methods) and immunohistochemistry (IHC) analysis revealed substantially higher vascular TfR expression in neonates that decreased at 1.5-3 months and became further reduced in 12-24 month-old adults (**Fig. S1b, Fig. 1b-c**) despite an increase in total vascular area by 24 months (**Fig. S1c**). We further validated these findings by using flow cytometry to isolate brain endothelial cells (BECs) from animals spanning 1.5 to 28 months of age. This method was not technically feasible for neonatal animals due to insufficient sample volume. Consistent with the IHC results, evaluation of TfR protein expression in CD31^+^ FACS-isolated BEC lysate revealed the highest TfR expression in 1.5 and 3 month-old mice, with a subsequent reduction at 16 months and a further reduction at 28 months (**Fig. 1d**).

**Fig. 1.**
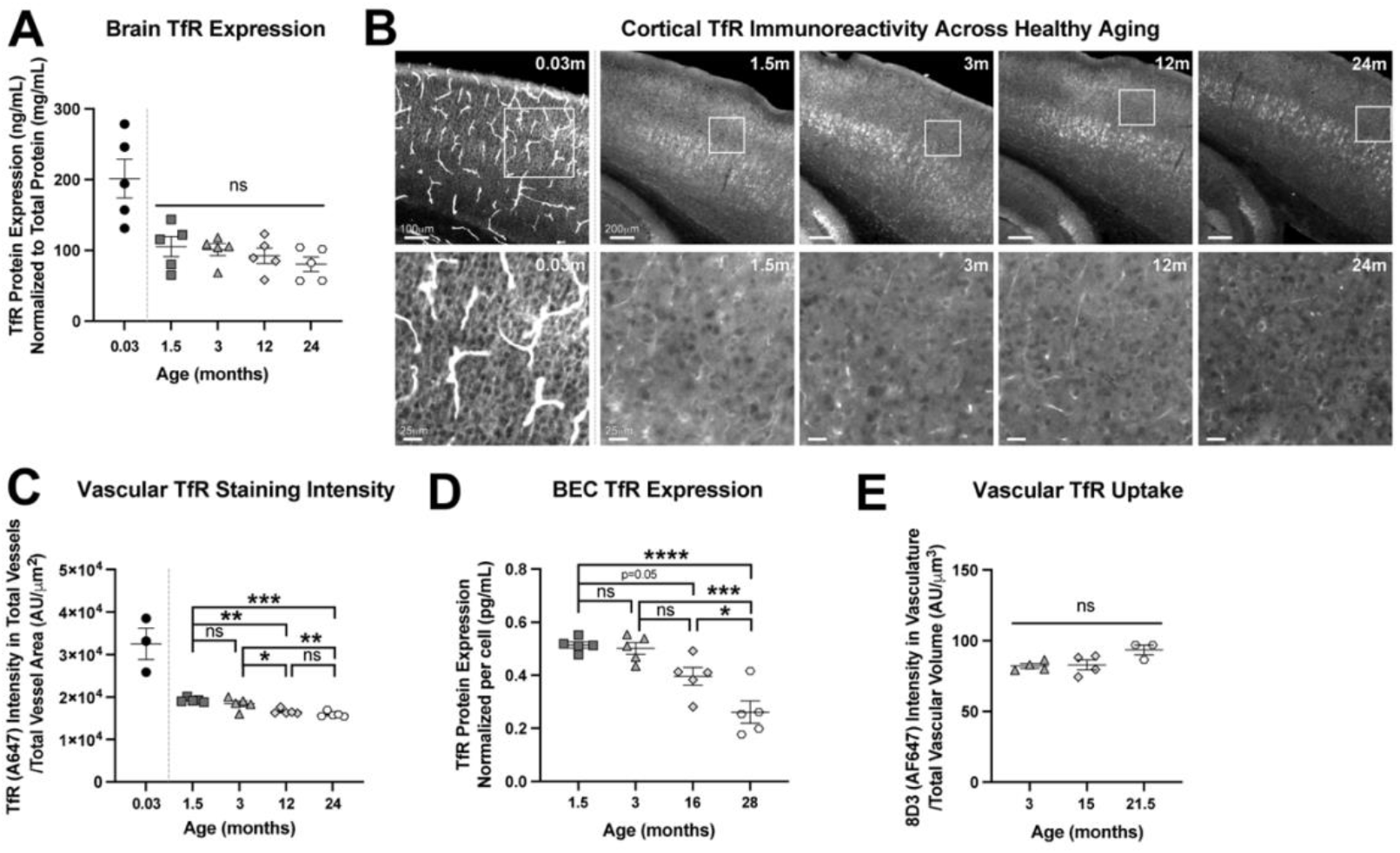
Vascular TfR expression is highest in young mice and vascular uptake of TfR antibodies remains consistent throughout adulthood. (**A**) TfR protein expression across ages quantified by ELISA and normalized to total protein in bulk brain lysate of TfR^mu/hu^ KI mice. Dashed line represents the exclusion of neonates from statistical analysis to allow for the detection of more subtle changes in TfR expression across aging. (**B**) Representative images of TfR immunoreactivity in the cortex (top row) of TfR^mu/hu^ KI mice with zoomed insets from cortical layer IV below (top row - neonatal (0.03 month) scale bar 100mm, juvenile and adult (1.5-24 months) scale bar 200mm; bottom row - all scale bars 25mm). Due to high TfR signal intensity, the brightness of the neonatal images was set to a separate threshold when compared to other age groups. (**C**) Mean TfR intensity in IHC-segmented vasculature across age groups in sagittal brain sections of TfR^mu/hu^ KI mice. (**D**) TfR expression per sorted CD31^+^ BEC from TfR^mu/hu^ KI mice. (**E**) Quantification of mean vascular uptake of AF647-labeled 8D3 (anti-murine TfR) in tissue-cleared hemibrains of WT mice. (**A, C-E**) one-way ANOVA, Tukey’s post hoc. (**A, C-E**) Graphs display mean ± SEM, n=3-5. * p<0.05, ** p<0.01, *** p<0.001, with only relevant comparisons indicated on graph; see Tables S1-2 for exact p values of all group comparisons. Images shown in (**B**) represent an n=3-5 animals per age group, with an n=2 brain sections per animal.

The luminal cell surface (i.e., blood-facing) pool of BEC TfR is the most relevant for the transport of systemically administered TfR-targeted therapeutics into the brain. To further understand whether the age-dependent reduction of TfR in the vasculature applies to this blood-accessible pool of TfR, we systemically dosed an Alexa Fluor 647 (AF647)-conjugated murine TfR antibody 8D3 (AF647-8D3) and evaluated vascular signal intensity within 1 hour post-dose in tissue-cleared wild-type (WT) mouse brains using light sheet fluorescence microscopy (LSFM). This time point was chosen because we previously demonstrated that systemically administered ATV^TfR^ immunoreactivity is predominantly localized to the vasculature a few hours post-dose (*14*). Additionally, 8D3 is expected to be mostly restricted to the brain vasculature due to its high-affinity, bivalent architecture (*35, 36*). Utilizing 8D3 in WT mice therefore allowed us to evaluate how age impacts the pool of blood-accessible vascular TfR. BEC binding and internalization of AF647-8D3 remained stable across adult aging in WT mice (**Fig. 1e, Fig. S1e**), despite an increase in total vascular volume with advancing age (**Fig. S1d**). Taken together, these data suggest that vascular TfR expression is reduced with age while vascular area and volume increase with age. Importantly, TfR-mediated uptake at the luminal surface of BECs appears to be functionally stable across age in adult mice.

### ATV^TfR^ transport across the BBB is not impacted by aging in adult mice

To understand whether the differences in total vascular versus luminal TfR expression functionally impact TfR-mediated antibody transport into the brain across adult aging, plasma and brain concentrations of two affinity variants of ATV^TfR^ with non-binding Fabs were evaluated in adult mice 24 hours after a single 25 mg/kg intravenous (i.v.) dose compared to a control huIgG with the same non-binding Fabs (ATV^TfR^-A: hTfR k_D_ = ∼1100 nM; ATV^TfR^-B, hTfR k_D_ = ∼100 nM). Due to the expected TfR-mediated clearance, both ATV^TfR^ molecules exhibited affinity-dependent reduction in plasma exposure compared to control huIgG (**Fig. 2a**). ATV^TfR^-A and ATV^TfR^-B achieved significantly higher brain exposure compared to control huIgG at all ages in an affinity-dependent manner (**Fig. 2b)**, consistent with previous reports (*37*). Brain exposure was modestly reduced at 24 months for the weaker affinity ATV^TfR^-A, and at both 10.5 and 24 months for the stronger affinity ATV^TfR^-B _(**Fig. 2b**)_. Importantly, the brain-to-plasma ratio of ATV^TfR^ exposure remained stable across age (**Fig. 2c**), suggesting that the reduction in the observed absolute brain concentration was predominantly driven by the age-dependent reduction in plasma exposure, rather than by altered transport capacity across the BBB.

**Fig. 2.**
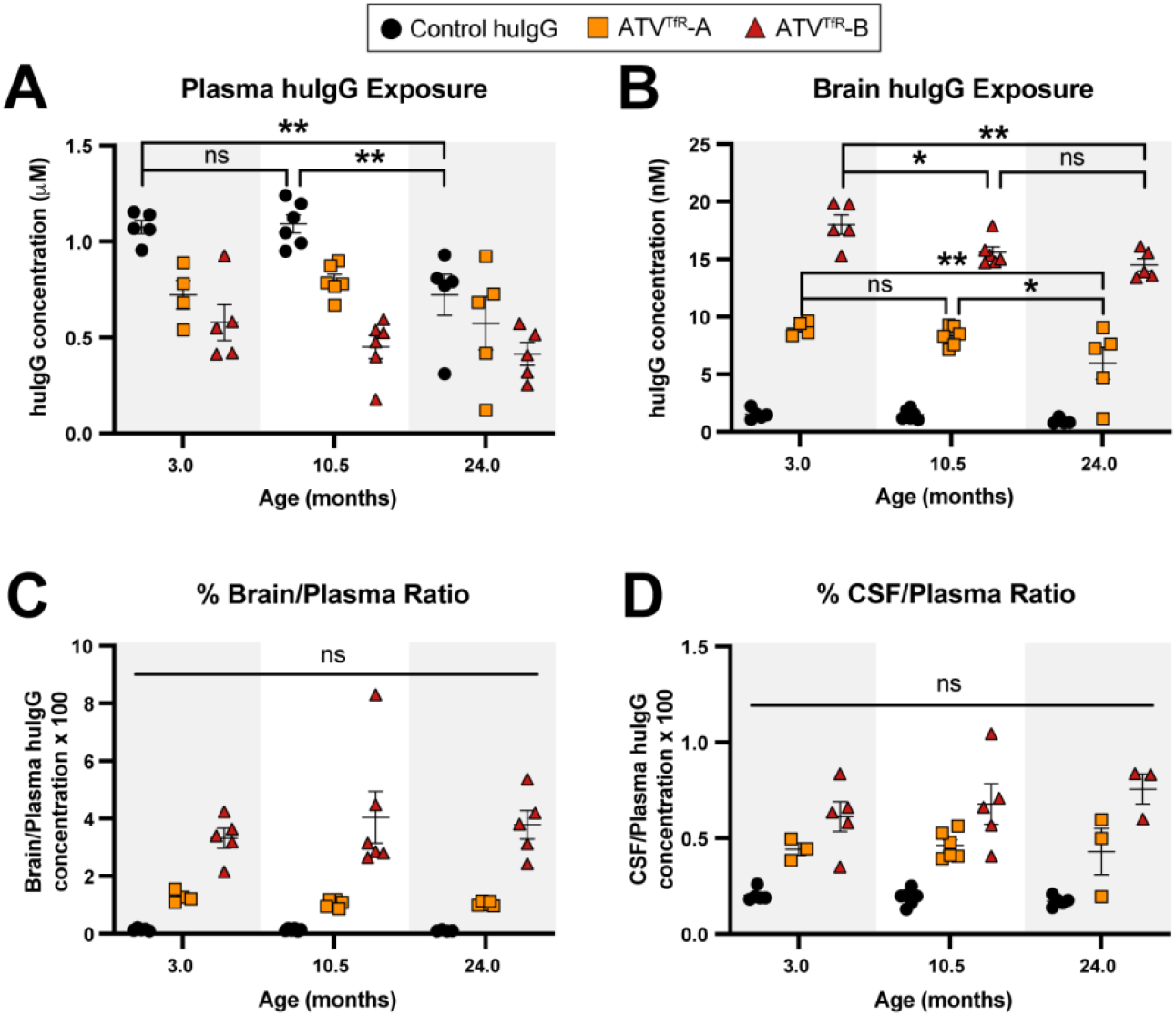
TfR-mediated transport of ATV^TfR^ across the BBB is not impacted by adult aging. (**A-B**) absolute huIgG concentrations of control huIgG or ATV^TfR^ 24 hours after a single 25mg/kg intravenous (i.v.) dose in plasma (A) and bulk brain lysate (B) of TfR^mu/hu^ KI mice. (**C-D**) Brain-to-plasma (C) and CSF-to-plasma (D) ratios of huIgG 24 hours after post-dose. (**A-D**) two-way ANOVA, Tukey’s post hoc, graphs display mean ± SEM, n=4-6. ** p<0.05, ** p <0.01, with only relevant comparisons indicated on graph; see Tables S1 and S3 for exact p values of all group comparisons. ATV^TfR^-A hTfR = ∼1100nM K_D_; ATV^TfR^-B = hTfR ∼100nM K_D_

Recent reports have also suggested that BBB permeability may be increased with age and contribute to an increase in non-specific transport into the CNS with age (*17*). However, we did not observe an increase in BBB permeability of exogenously administered control huIgG across 3-24 months (**Fig. S2b**). Similarly, there was no age-dependent change in CSF exposure or transport efficiency for either control huIgG or ATV^TfR^ molecules (**Fig. S2a, Fig. 2d**), suggesting that transport across the blood-CSF-barrier (BCSFB) was also not impacted by age. Our data therefore suggest that healthy adult aging between 3-24 months does not significantly impact the transport efficiency of TfR-targeting ATV molecules into the brain or CSF. Similarly, a lack of change in the brain-to-plasma ratio of exogenous IgG across 3-24 months of age also suggests that nonspecific IgG transport efficiency remains unchanged over this age range.

### ATV^TfR^ transport capacity is highest in young mice and maintained throughout healthy aging

We next evaluated time course brain uptake kinetics and compartmental biodistribution (i.e., isolated brain vascular and parenchymal fractions) across a broader age range and at a more therapeutically relevant ATV^TfR^ dose level to further explore TfR-mediated transport capacity. We focused on the ATV^TfR^-A variant and evaluated brain concentrations at 1, 4, and 7 days following a single 10 mg/kg dose. Given the elevated vascular TfR expression observed in neonates and 1.5 month-old juveniles **(Fig.1b-d)**, we included these two additional age groups to assess the possible impact of their increased TfR expression on functional transport over one week. We observed an age-dependent impact on plasma pharmacokinetics (PK) with ATV^TfR^-A, where clearance was modestly increased in juvenile 1.5 month-old mice as compared to adult mice (**Fig. 3a**). There was no difference in plasma clearance of control huIgG across 1.5-24 months. Neonates exhibited nearly 2-4-fold higher brain exposure of ATV^TfR^ at 1 day post-dose compared to all other age groups (**Fig. 3b**). The highest brain exposure occurred at 1 day post-dose for all age groups, suggesting that the kinetics of ATV^TfR^ uptake are consistent across ages. The brain-to-plasma ratio for ATV^TfR^ was over 1.5-fold higher in juvenile mice compared to the older age groups and nearly 9-19-fold higher than control huIgG depending on age (**Fig. 3c**, compared to **Fig. 2c**). These data suggest that TfR-mediated transport across the BBB and delivery of ATV^TfR^-A into the brain peak in early development and stabilize thereafter. Interestingly, brain exposure for control huIgG also peaked in neonates and remained consistent from the juvenile period through 24 months of age (**Fig. 3b, Fig. S2c**). CSF uptake kinetics and BCSFB transport of control huIgG remain consistent between 1.5 and 24 month-old animals (**Fig. S2d-f)**. The BBB of the early developing postnatal brain has been described as relatively impermeable to even small molecules (*38*). The increase in control huIgG brain exposure in neonatal animals may reflect slower CSF turnover (*38*) and/or increased IgG transit across the BCSFB to the CSF (and from there, to the brain (*39*)), consistent with higher CSF protein levels that have been reported in neonatal rodents (*40*) and human beings (*41*).

**Fig. 3.**
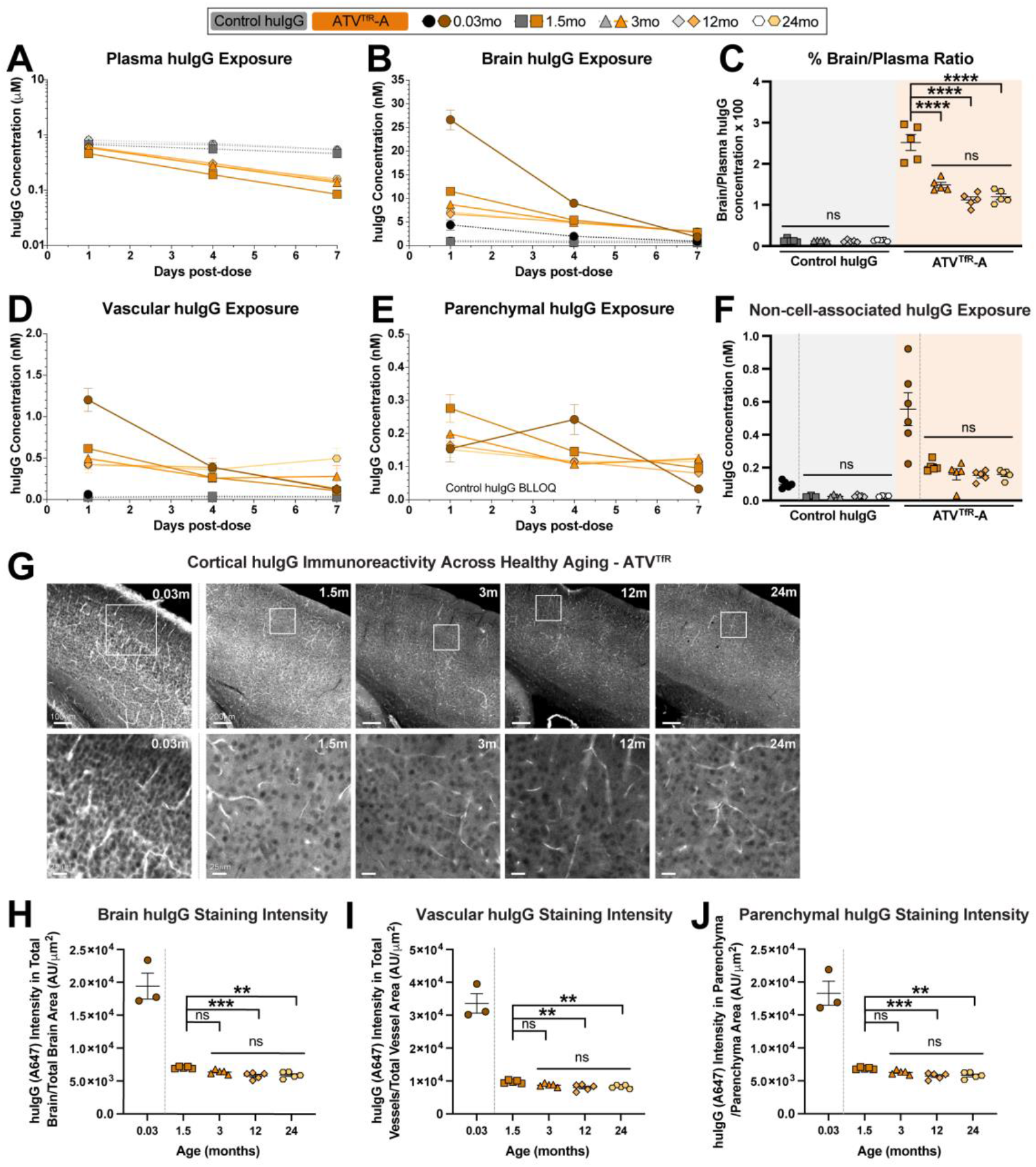
TfR-mediated transport capacity is highest in young mice and is maintained across healthy aging. TfR^mu/hu^ KI mice were administered a single 10 mg/kg i.v. dose of either control huIgG or ATV^TfR^-A. (**A-B**) absolute huIgG concentrations of control huIgG and ATV^TfR^-A in plasma (A) and bulk brain lysate (B) across 7 days. (**C**) Brain-to-plasma ratio of huIgG 24 hours post-dose. (**D-E**) absolute huIgG concentrations in vascular (D) and parenchymal (E) brain fractions separated by capillary depletion across 7 days. Any sample measurements below the lower limit of assay quantification are indicated by ‘BLLOQ’. (**F**) absolute huIgG concentration in the non-cell-associated fraction obtained by capillary depletion 24 hours post-dose. (**G**) Representative images of cortical huIgG immunoreactivity (top row) with zoomed insets from cortical layer IV below 24 hours post-dose (top row - neonatal (0.03 month) scale bar 100mm, juvenile and adult (1.5-24 months) scale bar 200mm; bottom row - all scale bars 25mm). Due to high huIgG signal intensity, the brightness of the neonatal images was set to a separate threshold when compared to other age groups. Dashed line indicates the exclusion of neonates from statistical analysis to allow for the detection of more subtle changes in huIgG immunoreactivity across adulthood. (**H-J**) Quantification of mean huIgG signal segmented by IHC 24 hours post-dose within total brain (H), vascular (I), and parenchymal (J) compartments in sagittal brain sections. (**A-B, D-E**) see mean results in Supplemental Table 1, (**C, F**) two-way ANOVA, Tukey’s post hoc, (**H-J**) one-way ANOVA, Tukey’s post hoc. (**A-F, H-J**) Graphs display mean ± SEM, n=4-6. * p<0.05, ** p<0.01, *** p <0.001, **** p <0.0001, with only relevant comparisons indicated on graph; see Tables S1 and S3 for exact p values of all group comparisons. Representative images are shown from n=5-6 animals/treatment groups, n=2-3 brain sections per animal.

Given the difference in TfR-mediated brain uptake in juvenile versus adult mice, we aimed to more fully understand whether transcytosis kinetics at the BBB differ with age. We therefore used brain capillary depletion to separate out vascular and parenchymal fractions in order to evaluate ATV^TfR^-A uptake across the 7 day time course. At 1 day post-dose, neonates exhibited over 2-fold higher ATV^TfR^-A exposure in the vasculature, whereas parenchymal exposure remained mostly similar across all ages (**Fig. 3d, e**).

Taken together, these results suggest that the elevated overall brain exposure measured in neonates 1 day post-dose is primarily driven by elevated vascular localization rather than enhanced parenchymal exposure. Interestingly, peak parenchymal ATV^TfR^-A exposure appeared to be delayed in neonates when compared to the other age groups (**Fig. 3e**). Lastly, to further evaluate whether age impacts delivery to the brain parenchyma, ATV^TfR^-A exposure in segmented vascular and non-vascular (i.e., parenchyma) compartments was quantified by IHC in sagittal brain sections at 1 day post-dose, corresponding to peak brain exposure (**Fig. 3g-j**). Consistent with ATV^TfR^-A exposure in whole brain and vascular lysates (Fig. 3b, d), ATV^TfR^-A exposure in the total brain (**Fig. 3g-h, Fig. S2g**) and segmented vasculature (**Fig. 3i**) was highest in neonates, with an age-dependent rank order that remained consistent throughout adulthood. Together, this data reinforces that TfR-mediated brain uptake of ATV^TfR^-A is highest during early development and adolescence, while the remainder of adulthood is characterized by stable brain uptake that remains significantly higher than that of control IgG.

Interestingly, our IHC analysis revealed the highest parenchymal ATV^TfR^-A exposure in neonates (**Fig. 3j**), which is inconsistent with what was observed when measuring huIgG in tissue lysate (Figure 3e). We hypothesized that a significant fraction of the parenchymal ATV^TfR^-A exposure detected by IHC in neonatal mice could be non-cell-associated (i.e., extracellular), as huIgG measurements in tissue lysates primarily reflect cell-associated huIgG. To test this hypothesis, we separated whole brain tissue into cell-associated (i.e., both vascular and parenchymal) and non-cell-associated fractions (see Materials and Methods). Elevated levels of non-cell associated ATV^TfR^-A exposure were indeed observed in neonates when compared to older mice (**Fig. 3f**). Importantly, this finding is consistent with the low parenchymal exposure detected in neonatal tissue lysate as compared to IHC, indicating that a significant amount of ATV^TfR^-A was indeed present in the non-cell associated fraction, likely resulting from reduced parenchymal cell TfR expression at this early postnatal age (*42*).

### 5xFAD disease progression does not significantly impact TfR-mediated transport of ATV^TfR^

There have been few studies directly evaluating whether neurodegenerative disease impacts TfR protein expression at the BBB and/or TfR-mediated transport capacity. This is particularly relevant in AD, where vascular amyloid accumulation could potentially impact BBB physiology and transport. To address this, we evaluated TfR expression and brain uptake of ATV^TfR^-A in TfR^mu/hu^ KI crossed to the 5xFAD amyloid deposition mouse model (*43, 44*) at 1.5, 3, and 10.5 months of age. These ages were chosen to represent advancing stages of amyloid beta plaque burden (**Fig. 4a**). Total brain TfR expression remained unchanged across age and genotype (**Fig. 4b**), although some differences in sorted BEC TfR expression were observed (**Fig. 4c**). To determine whether TfR-mediated transport was altered in 5xFAD mice across ages, plasma, brain, and CSF exposures were assessed 24 hours following a single systemic dose of ATV^TfR^-A. Plasma exposures were similar between TfR^mu/hu^ KI and 5xFAD; TfR^mu/hu^ KI mice across ages (**Fig. 4d**). Importantly, disease progression had minimal impact on whole brain, vascular, or parenchymal ATV^TfR^-A uptake as well as CSF exposure, as huIgG levels were similar between TfR^mu/hu^ KI and TfR^mu/hu^; 5xFAD KI mice at each age (**Fig. 4e-h**). Similarly, brain-to-plasma and CSF-to-plasma ratios were unchanged between TfR^mu/hu^ KI and 5xFAD; TfR^mu/hu^ KI mice for control huIgG and ATV^TfR^-A (**Fig. 4i-j**). The reduction in brain-to-plasma ratio between 1.5 and 3-10.5 months of age is consistent with what was observed in an earlier study comparing healthy juveniles and adult mice (Fig. 3c).

**Fig. 4.**
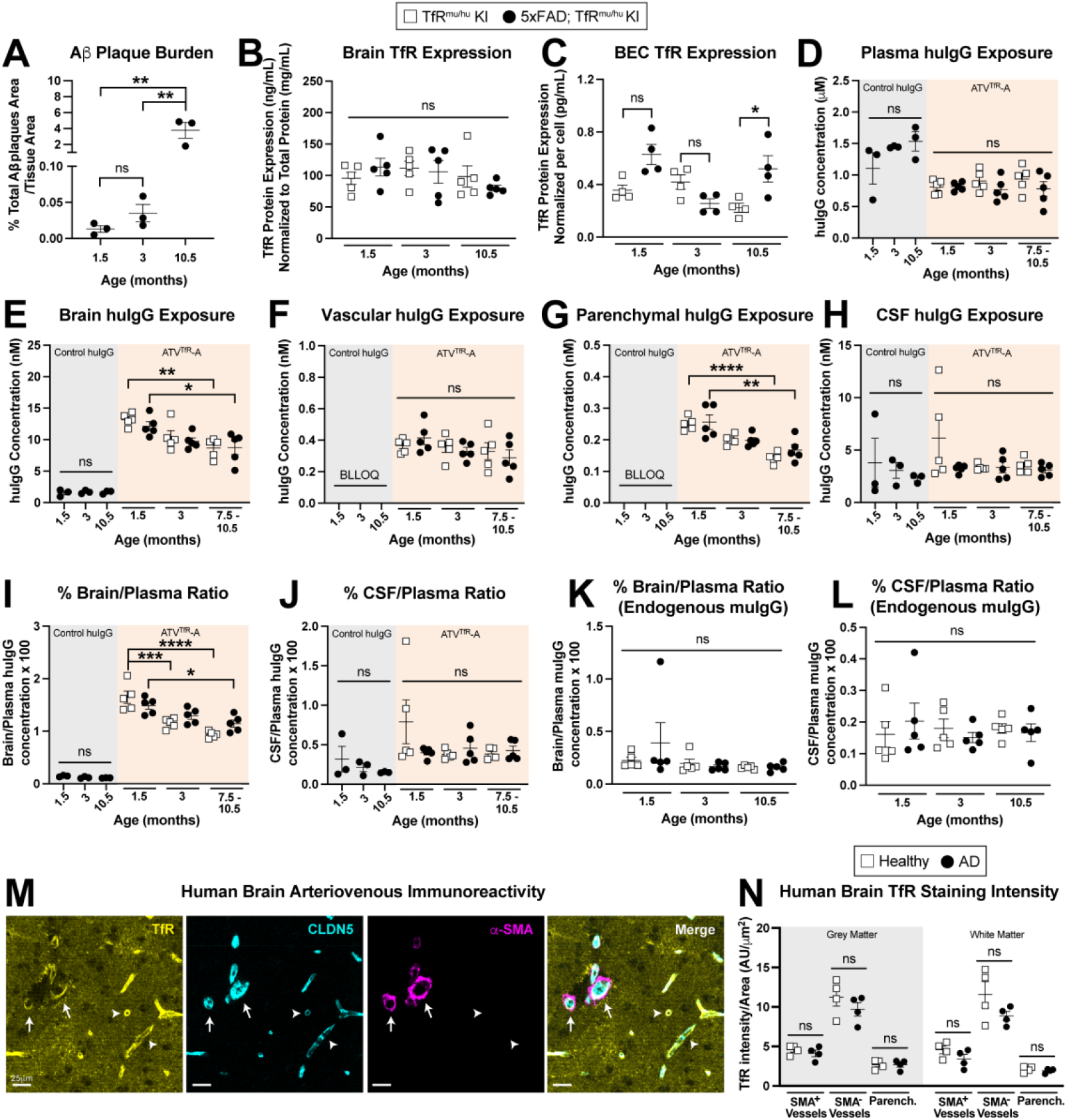
Disease progression does not impact ATV^TfR^-mediated transport across the BBB in 5xFAD;TfR^mu/hu^ KI mice or vascular TfR expression in AD patient brains. (**A**) Mean amyloid beta plaque burden measured by IHC in sagittal brain sections of 5xFAD;TfR^mu/hu^ KI mice across ages. (**B-C**) TfR protein expression in bulk brain (B) and sorted CD31^+^ BECs (C) lysate from 5xFAD;TfR^mu/hu^ KI mice (note that n=2 animals were pooled per individual data point shown in (C)). (**D-H**) absolute huIgG concentrations of control huIgG or ATV^TfR^-A 24 hours after a single 10 mg/kg i.v. dose in plasma (D), bulk brain lysate (E), isolated vascular (F) and parenchymal brain fractions (G) separated by capillary depletion, and CSF (H) of 5xFAD;TfR^mu/hu^ KI mice. (**I-J**) Brain-to-plasma (I) and CSF-to-plasma (J) ratios of huIgG 24 hours after a single 10 mg/kg i.v. dose of control huIgG or ATV^TfR^-A. (**K-L**) Brain-to-plasma (K) and CSF-to-plasma (L) ratios of endogenous muIgG across ages. (**M**) Representative image of the grey matter of the frontal cortex in an 87-year-old human AD patient brain qualitatively illustrating the expected zonal pattern of TfR protein along the arteriovenous axis (TfR, CLDN5, and a-SMA; scale bar 25mm). Arrows identify large caliber (≥20mm) arteries/arterioles, whereas arrowheads identify smaller caliber microvessels (<10mm). (**N**) Quantification of TfR protein intensity within a-SMA^+^ arteries, a-SMA^-^ vessels, and brain parenchyma in the grey and white matter of control (81-86-year-old) and age-matched AD (80-96-year-old) patient brains. (**A**) one-way ANOVA, Tukey’s post hoc, (**B-L**) two-way ANOVA, Tukey’s post hoc. (**A-L**) Graphs display mean ± SEM, n=3-5; see Table S5 for human patient demographics. * p<0.05, ** p<0.01, *** p<0.001, **** p<0.0001, with only relevant comparisons indicated on graph; see Table S4 for exact p values of all group comparisons.

It has been suggested that AD pathology may lead to an increase in CNS barrier permeability (*19, 26*). We observed no age-dependent difference in brain exposure of exogenously administered control IgG (**Fig. 3e, Fig. 3i**). Endogenous IgG was found to be increased in plasma, brain, and CSF in an age-dependent manner (**Fig. S3a-c**), but the brain-to-plasma and CSF-to-plasma ratios remained stable across age and disease progression (**Fig. 4k-l**). These data suggest that increased amyloid burden in this disease model did not compromise the BBB or BCSFB integrity sufficiently to permit increased non-specific leakage of endogenous IgG or exogenously administered huIgG. Together, these data suggest that the progression of pathology in the 5xFAD mouse model has little to no effect on the transport of IgG into brain or CSF and that transport of ATV^TfR^ into the brain does not significantly differ between 5xFAD and normal mice across 1.5-10.5 months.

### TfR expression along the human brain arteriovenous axis is unaffected by Alzheimer’s disease

We undertook additional studies in human tissue to better appreciate TfR expression patterns between AD and control brains. TfR mRNA has been reported in transcriptomics studies (*20, 22, 45-47*) to be highest in human capillary/venous BECs, with lower levels in arterial BECs, an example of arteriovenous ‘zonation’ (*48*). However, an evaluation of TfR protein zonation along the human arteriovenous axis has not yet to our knowledge been reported. We utilized multiplex IHC on cortical brain tissues of male and female AD patients and healthy age-matched controls to compare vascular TfR zonation in AD and healthy, age-matched control brains. Claudin 5 (CLDN5) was used as a general vascular marker and alpha smooth muscle actin (a-SMA), which is largely absent on capillaries and veins, was used as a marker to identify larger caliber arteries and arterioles. In both AD (**Fig. 4m**) and healthy age-matched control (**Fig. S4a**) brains, CLDN5^+^ microvessels (<10mm) that were devoid of a-SMA immunoreactivity exhibited high TfR expression; in contrast, larger caliber vessels (≥20mm; putative arterioles/arteries) that were CLDN5^+^ and a-SMA^+^ exhibited low TfR expression. TfR expression (**Fig. S4b**) was quantified in segmented vasculature (a-SMA^+^, TfR^+^ arteries or a-SMA^-^, TfR^+^ vessels) and parenchyma of the grey and white matter (**Fig. S4c**). Vascular and parenchymal TfR expression in both the grey and white matter (**Fig. 4n**) were unaffected by AD disease state when compared to healthy age-matched controls. Our findings demonstrate arteriovenous zonation of TfR protein in human brains and suggest that TfR protein expression patterns are similar between AD and healthy control brain tissue.

## Discussion

Our study produced several key findings. First, we demonstrate that vascular TfR expression is significantly elevated in neonatal compared to adult mice, leading to significantly increased neonatal brain exposure of a TfR-targeted ATV. Second, we confirm vascular TfR expression in BECs modestly declines in adult mice, most evident when comparing younger 1.5 month-old mice to mice 24 months or older. This TfR reduction is accompanied by a modest age-dependent decrease in ATV^TfR^ brain exposure and brain-to-plasma ratio, suggesting a reduction in transport efficiency at the BBB in adult mice when compared to juvenile mice. Importantly, our study demonstrates stable vascular TfR uptake and ATV^TfR^ brain-to-plasma ratios in mice 3 months and older in both healthy aging and in a mouse model of AD. Finally, we provide new evidence to show that TfR levels are largely maintained between healthy and diseased brains from 5xFAD mice and AD patients. These results provide among the most comprehensive data to date on TfR protein expression and TfR-mediated transport into the brain across mouse development with potential translational significance for the prediction of TfR-mediated transport in humans across a wide age range.

Our finding demonstrating elevated vascular TfR expression in neonatal mice is consistent with previous reports suggesting the upregulation of TfR during early development (*42, 47, 49, 50*) and its reduction in aged animals (*17, 18, 24, 31, 51*). Elevated TfR during the neonatal period was accompanied by markedly increased brain concentrations of ATV^TfR^. This finding may hold special significance for the future application of TfR-targeted enzyme replacement therapies (e.g. the enzyme transport vehicle (*52*)) when administered to younger children and infants in the treatment of neuronopathic lysosomal storage disorders. Another important inference from our study was that the reduction in vascular TfR during adulthood did not appear to functionally impact TfR-mediated brain uptake of IgG in mice across 3-24 months of age. Although the age-dependent changes in vascular TfR expression we observed were generally consistent with previous reports (*17, 18, 24, 31*), our study is the first to demonstrate that TfR-mediated brain delivery at a therapeutically relevant dose level is not significantly impacted by this reduction. Because TfR undergoes rapid ligand-independent recycling kinetics, it is possible that TfR-mediated transport may be less susceptible to changes in absolute TfR expression (i.e., reduced copy number of TfR may not functionally impact transport if fast recycling is able to quickly traffic molecules intracellularly) (*53, 54*). Taken together, our results suggest that sufficient TfR is expressed at the luminal BEC surface to effectively maintain transport into the brain across aging in adult animals. Future evaluation will be needed to determine whether aging and disease may impact the expression and transport of other BBB targets currently being studied for CNS drug delivery that have different receptor trafficking kinetics (*55-57*).

We speculate that the subtle reduction in BEC TfR we observed across healthy adult aging could reflect a regulatory mechanism for maintaining iron homeostasis in the brain. Iron is elevated in the aged brain (*58-60*) and strongly associated with AD pathology in mouse models (*61-64*) and humans (*65*). The elevated TfR we observed in sorted BECs from 5xFAD mouse brain compared to controls at 10.5 months may indicate regulatory perturbation. ATV^TfR^ levels were unchanged across 1.5 to 10.5 months in whole brain, vascular, and parenchymal fractions between 5xFAD mice and age-matched controls, irrespective of age- or disease-dependent changes in vascular TfR expression and the significantly increased amyloid burden we observed in the 10.5 month-old animals. Prior studies have also demonstrated preserved brain exposures for TfR-targeted antibodies compared to healthy age-matched controls in multiple other AD mouse models (*29, 30*).

In summary, our data provides new mechanistic insights into the impact that changes in TfR expression may exert on TfR-mediated transport in healthy aging and a mouse model of AD. Future studies will be needed to evaluate how vascular TfR expression and TfR-mediated transport may be impacted by other age-related neurodegenerative CNS diseases that may benefit from this delivery platform. Our finding of substantially increased ATV^TfR^ brain exposure during early development also highlights the potential benefit that TfR-targeted platforms may have in the treatment of neurological diseases that impact the young (e.g., lysosomal disorders). Both preclinical studies (*52, 66*) and on-going clinical trials (*27*) utilizing the TV^TfR^ platform for enzyme delivery have thus far yielded promising results. Nevertheless, the addition of other therapeutic payloads (e.g. targeted Fabs) to the TV^TfR^ platform may alter transport properties. Understanding the influence of developmental stage and disease effects will therefore likely require further evaluation on a case-by-case basis. This topic may take on increasing importance as TfR-based drugs continue to emerge as a promising class of neurotherapeutics.

## Materials and Methods

### Animal care and dosing

All procedures in animals were performed with adherence to ethical regulations and protocols approved by Denali Therapeutics Institutional Animal Care and Use Committee. Mice were housed under a 12 hour light/dark cycle (lights on at 7 AM, lights off at 7 PM) with *ad libitum* access to water and a standard chow (Labdiet, #5LG4) unless otherwise indicated. Whenever possible, animals were group housed of up to 5 sex-matched animals/cage. Both male and female TfR^mu/hu^ KI mice (*14*), transgenic 5xFAD (familial AD) mice (*43, 44*) crossed to TfR^mu/hu^ KI mice (5xFAD; TfR^mu/hu^ KI), and wild-type (WT) mice of C57BL/6J background (The Jackson Laboratory) were used between 0.03 and 28 months of age with approximately equal distribution between treatment groups based on sex and age. Although neonatal dosing differed from that of juvenile and adult mice (see Supplemental Information for detailed *in vivo* dosing methods across age), all animals were intravenously (i.v) administered either control huIgG or ATV^TfR^ with the same anti-DNP (*67*) non-binding Fabs at a volume of 10 mg/kg body weight.

### Brain endothelial cell isolation

Olfactory bulbs and cerebellum were grossly dissected from fresh, PBS-perfused brains and the remaining tissue was mechanically dissociated on ice with a razor and collected into a 15mL conical containing 6mL cold HBSS. Following centrifugation at 300G for 3 minutes at 4°C, the supernatant was aspirated and brain samples proceeded to enzymatic dissociation (Miltenyi 130-092-628) according to the manufacturer’s protocol. All samples were triturated 10x between the addition of enzymes and upon completion, samples were washed through a 100µm filter with 20mL ice-cold DPBS and centrifuged at 300G for 10 minutes at 4°C. The supernatant was aspirated, and the pellet resuspended in 45mL ice-cold 0.9M sucrose (Sigma 84097) and centrifuged at 850G for 30 minutes at 4°C (slow breaks). The resulting supernatant was aspirated, and the cell pellet was washed with DPBS at 1,600RPM for 10 minutes at 4°C. The resulting supernatant was aspirated, and cell pellet was resuspended in FACS buffer (PBS + 1% BSA + 2mM EDTA) and washed again at 1,600RPM for 10 minutes at 4°C. Cells were then blocked with 1:10 FC block (Miltenyi 130-092-575) for 15 minutes at 4°C, followed by a 30 minute incubation of a FACS antibody mix (Ghost Dye-v450 (Tonbo 13-0863-T500; 1:1,000), CD45-PE-Cy7 (Biolegend 103114; 1:200), CD31-PE-CF594 (BD 563616; 1:75), CD11b-APC-Cy7 (Tonbo 25-01120-U100; 1:100), Thy1-A488 (R&D FAB7335G; 1:100), ACSA2-PE (Miltenyi 130-123-284; 1:100), O-1-APC (Invitrogen 50-6506-80; 1:100)) at 4°C protected from light. Stained cells were then washed with FACS buffer and centrifuged at 1,600RPM for 10 minutes at 4°C. Following the aspiration of supernatant, cell pellets were resuspended in 0.5mL FACS buffer and 25,000 BECs were sorted (>98% purity of Ghost Dye^-^, CD45^-^, CD31^+^, CD11b^-^, Thy1^-^, ACSA2^-^, O-1^-^) on the BD FACS ARIA III into standard sorting collection buffer (PBS + 10% FBS (VWR 89510-188)). BECs were then centrifuged at 10,000G for 10 minutes at 4°C, the supernatant was aspirated, and the remaining cell pellet was lysed at a final concentration of 1.25×10^6^ cells/mL in lysis buffer (1% NP40-PBS containing protease and phosphatase inhibitors (Roche)) and stored at -20°C until further analysis.

### Immunohistochemistry (IHC), vascular labeling, and microscopy

Mouse and human fixed brain sections underwent independent IHC staining (see Supplemental Information for IHC protocols and microscopy analysis); both tissues were generally photobleached to reduce tissue autofluorescence, blocked, and stained with a mixture of primary antibodies that were detected either by secondary antibody staining (visualized on a Zeiss Axioscan.Z1 microscope) or by a PhenoCycler Fusion antibody barcode system (visualized on a Sony IMX421-based camera), respectively. Mouse 3D hemibrain vascular labeling was performed *in vivo* (see Supporting Information for the vascular labeling protocol and microscopy analysis), where animals were i.v. dosed with a fluorescently conjugated TfR antibody, perfused with fluorescently conjugated lectin, and then fixed with 4% paraformaldehyde (PFA). Dissected, intact PFA-fixed hemibrains then proceeded to F-DISCO tissue clearing and the vascular labeling was visualized on a Miltenyi Ultramicroscope Blaze Lightsheet microscope.

### Statistical Analysis

As indicated in figure legends, all data is expressed as mean ± SEM and all statistical analyses were performed in GraphPad Prism v10.3.1. Analyses were done using either one- or two-way ANOVA, and as indicated in figure legends, the criterion for differences to be considered significant was p<0.05. The neonatal data skewed the data distribution for an initial parametric analysis across all ages (Fig. S1a-b, Fig. S2e, j, and Tables S2-3), and thus, were excluded from a secondary ANOVA analysis to allow for parametric analysis on normal data distribution across juveniles and adults (Fig.1 a-c, Fig. 3f-j, and Tables S2-3). Any sample that was missing due to it being below the lower limit of quantification (BLLOQ) was imputed as the lower limit of quantification for analysis purposes. Tables S2-4 show the results for all one- and two-way ANOVA models, with additional Tukey post hoc analyses to determine pairwise differences between any two groups of each individual ANOVA model.

## Acknowledgments

The authors would like to thank Drs. Arash Moshkforoush and Audrey Gill for their intellectual assistance throughout this study. The authors would also like to thank Butch Benitez and Kevin Rebadulla for technical assistance in animal colony maintenance across all aging studies. Lastly, the authors would like to thank Najiba Mammadova, Oliver Braubach, Jasmine Singh, Katrina Evans, and Bassem Ben Cheikh at Akoya for their invaluable guidance on the human CODEX imaging and analysis.

## Funding

The financial support for this research was provided by Denali Therapeutics.

## Author contributions

Conceptualization: VOT, MEP, JWL, RJW, RGT, YJYZ

Methodology & Data Collection: VOT, MEP, JCD, DH, DJ, EKL, LS, IB, RC, KSC, JC, TKE, NK, KJL, AWL, HNN, ESO, ER, HS, MT, RKT

Formal Analysis & Investigation: VOT, MEP, DC, JCD, DH, DJ, EKL, KH, MEC

Writing—original draft: VOT

Writing—review & editing: all authors commented read, edited, and approved the final manuscript

Supervision: RGT, YJYZ

## Competing interests

All authors are current or past employees of Denali Therapeutics.

## Data and materials availability

All data are available in the main text and/or the supplementary materials.

## Supplemental Materials

Please see Supplemental Materials for additional methods (tissue collection, tissue processing, IHC, vascular labeling, and microscopy, microscopy image analysis, capillary depletion, TfR MSD, ATV^TfR^ and antibody generation, and IgG quantification), figures, and detailed tables describing statistical comparisons per figure and figure panel.

## Notes

### Competing Interest Statement

All authors were paid employees of Denali Therapeutics Inc. during the conduct of the study.

