## Supplemental Materials for "Transferrin receptor-mediated transport at the blood-brain barrier is elevated during early development but maintained across adult aging"

Vanessa O. Torres *et al.*

**Supplemental Materials and Methods:**

***In vivo* dosing:** Prior to handling neonatal animals, cage bedding was rubbed onto the gloves to transfer the home cage scent. The neonates were gently picked up from the nest and placed onto a weigh boat that contained nesting material from the home cage. The neonates were then individually administered the desired dosing solution (either control huIgG or $ATV^{TfR}$ with the same anti-DNP (*67*) non-binding Fabs) via jugular vein at a volume of 10mg/kg with needle bevel up and a cranial to caudal direction (BD Precision Glide needle, 30G x 1/2in, Hamilton syringe (80901) 50µL). The desired dosing solution was dispensed slowly, with a successful injection visually confirmed by clear fluid passing through the jugular vein. The neonate was then returned to the home cage and carried by the dam back to the nest. Each dosing solution had its own syringe and needle, and the syringe was refilled between neonates. For juvenile and adult animals, the desired dosing solution was administered via lateral intravenous (i.v.) tail vein injections at a volume of 10 mg/kg body weight. Animals were warmed under a heat lamp for ~5 minutes to dilate the veins and syringes (28.5G x 0.5ml Insulin Syringe, 26026) were carefully prepared with no air bubbles. Following vein dilation, animals were individually placed into a restrainer to prevent movement. The tail was subsequently cleaned with an alcohol wipe and the needle was inserted with the bevel up into the vein towards the direction of the animal’s head. Following a slow injection of the desired dosing solution, the needle was slowly removed from the tail vein and a gentle compression was applied at injection site until bleeding stopped.

**Tissue collection:** Postnatal day 2-6 neonates were placed on wet ice until deeply anesthetized and then cardiac perfused under a dissecting microscope using a 27G x 3/4in scalp vein set with ice cold PBS for 2-3 minutes at 10RPM. In contrast, postnatal day 7-10 neonates were deeply anesthetized via intraperitoneal (i.p.) injection with 2.5% tribromoethanol and then cardiac perfused using a 21G x 3/4in winged infusion set with ice cold PBS for 2-3 minutes at 15RPM. Juvenile and adult animals were humanely anesthetized with 2.5% tribromoethanol via intraperitoneal (i.p.) injection and transcardially perfused for 3-5 minutes at 5 mL/min with ice-cold PBS. At 24 hours, 4 days, and/or 7 days, juvenile and adult CSF extraction was achieved via cisterna magna puncture followed by centrifugation at 12,700RPM for 7 minutes at 4°C. The supernatant was then collected and also stored at -80°C until further analysis. Next, juvenile and adult plasma extraction was achieved by collecting whole blood via cardiac puncture into EDTA-coated tubes followed by centrifugation at 12,700RPM for 7 minutes at 4°C. The top plasma layer was then collected and stored at -80°C until further analysis. Lastly, brain tissue across all ages were harvested and dissected for desired endpoints (homogenization and IHC), weighed, and either frozen on dry ice and stored at -80°C until future tissue homogenization, or immersion fixed with 4% PFA for 24 hours and stored at 4°C until further IHC processing.

**Tissue processing:** Bulk brain tissue lysates were generated by homogenizing flash-frozen brain tissue in a 10x tissue weight volume of lysis buffer (1% NP40-PBS containing protease and phosphatase inhibitors (Roche)) with a 3mm tungsten bead for 6 minutes at 27Hz using a Qiagen TissueLyser II. The homogenate was spun down at 14,000G for 10 minutes at 4°C and the bulk brain lysate was transferred to a clean tube and stored at -80°C until further analysis. For IHC, brains were transferred to PBS + 0.01% sodium azide after having undergone the 4% PFA immersion fixation and were then shipped to Neuroscience Associates for sectioning (40µm-thick sections collected in a series of 12). Up to 20 hemibrains were embedded into a single gelatin block and sectioned via a microtome in the sagittal plane using MultiBrain Technology. Brain sections were collected and stored in antigen preservation solution (50% PBS: 50% ethlyene glycol + 1% PVP) at -20°C until staining.

**Immunohistochemistry (IHC), vascular labeling, and microscopy**:
**2D mouse IHC.** All NSA blocks of 40µm-thick, '2D' sagittal brain sections were subjected to an adapted photobleaching protocol (*68*) where tissues were exposed to broad-spectrum LED lights for 24-48 hours (ex/Aibecy A4 Ultra Bright 25,000 Lux LED Light Box-Tracing Pad) in 4.5% (w/v) $H_2O_2$ and 20mM NaOH in PBS to reduce as much age-related autofluorescence as possible prior to staining. Following photobleaching, tissues designated for TfR staining were first subjected to a series of 3-5 minutes gradient methanol dehydration (20% in $dH_2O$, followed by 40%, followed by 60%) and rehydration (60%, followed by 40%, followed by 20%) at room temperature to further fix the brain sections for optimal TfR staining. All tissues were then blocked in 5% BSA + 0.3% Triton X-100, followed by overnight staining with fluorescent primary antibodies (Ab plaques (mouse α-biotin Aβ (IBL America 10326)), vessels (goat α-CD31 (R&D AF3628) + goat α-podocalyxn (R&D AF1556)), neurons (chicken α-NeuN (Genetex gtx00837), and either TfR (rabbit α-TfR (Abcam ab214039)) or huIgG (A647-huIgG (Jackson Labs 709-606-149))) diluted in 1% BSA + 0.3% Triton X-100. Tissues were then washed with 0.05% PBS-TritonX and stained with secondary antibodies (α-biotin-Dy555 (Invitrogen 84606), α-goat-AF568 (Invitrogen A11057) or -AF488 (Invitrogen A11055), α-chicken-AF488 (Invitrogen A78948), and/or α-rabbit-AF647 (Invitrogen A31573), respectively), counterstained with DAPI, and mounted in Prolong glass (Thermofisher P36984). All NSA section blocks of '2D' sagittal brain sections were imaged on the Zeiss Axioscan.Z1 (20x objective, NA 0.8) microscope using Zeiss Zen 3.7 software. Lamp powers and exposure times were held constant across all images collected for each full set of fluorescently stained NSA section blocks from individual experiments/imaging sessions.

**3D mouse vascular labeling.** Female C57BL/6J WT mice (3-4-, 15-, and 21.5-month-old mice; Jackson Laboratory Strain 664) were used for the vascular labeling tissue clearing study. Animals first received a single 2.5 mg/kg dose of AlexaFluor 647-conjugated anti-TfR antibody (AF647-8D3 NB100-64979AF647, Novus) via tail vein injection. Following a 30 minute incubation post i.v. dosing of anti-TfR, animals were deeply anesthetized with 2.5% avertin (i.p.) and transcardially perfused with freshly prepared, ice-cold PBS containing 6 mg/L of conjugated lectin (LEL tomato lectin DyLight 594, L32471, ThermoFisher) for 6 minutes with a peristaltic pump at a rate of 2.5 mL/min. Animals were then immediately perfused with ice-cold PBS for 8 minutes, followed by room temperature 4% paraformaldehyde (PFA; diluted in PBS, 15714-S, Electron Microscopy Sciences) for 22 minutes. Brains were dissected and immersion fixed in 4% PFA overnight at 4°C before moving onto FDISCO-based tissue clearing methods (*69*). All steps were performed at 4°C with gentle agitation unless otherwise specified.

Following overnight 4% PFA immersion fixation, brains were dissected into two hemibrains along the sagittal midline. Hemibrains were washed with PBS for 2 hours followed by a 3-day wash with fresh PBS. After washing, hemibrains were dehydrated in a dilution series of Tetrahydrofuran (THF; Sigma-Aldrich 186562)/$dH_2O$. Hemibrains were first incubated in 50% THF/$dH_2O$ for 3 hours, followed by 70% THF/$dH_2O$ for 3 hours, and then incubated overnight in 80% THF/$dH_2O$. Following the overnight incubation, hemibrains were incubated overnight with 100% THF in a completely filled and sealed glass vial, followed by a 2-day similar incubation in fresh 100% THF. Following tissue dehydration, hemibrains were cleared for 3 hours at RT in pre-chilled dichloromethane (DCM; Sigma Aldrich 270997) until the completely sank in the glass vial, then were transferred to completely filled glass vials of RT dibenzyl ether (DBE; Sigma Aldrich 108014) to incubate 3 days in the dark at RT (undisturbed). Following the 3 days, hemibrains were switched to fresh Ethyl Cinnamate (ECi; Sigma Aldrich 112372) daily for 2 days and proceeded to imaging on the start of the 3$^{rd}$ day. Using the Miltenyi Ultramicroscope Blaze Lightsheet (imspector Pro v7.6.3 acquisition software), hemibrains were imaged in ECi with the LVBT 4x objective at 1x magnification, with a pixel size of 2.71 in X, Y, and Z planes. Anti-TfR-AF647 was imaged at 60% power (Lesos Beamcombiner) at 640nm excitation/680nm emission. Lectin-DY594 was imaged at 13% power at 561nm excitation/620 emission. Both left and right light sheets were activated at 100% sheet width during acquisition, creating a 3.86μM sheet, with a NA of 0.19.

**CODEX multiplexed IHC of human tissue.** Human frontal cortex brain tissue from 4 donors diagnosed with Alzheimer's Disease (AD$^+$, ages 80-96, all female) and from generally age-matched control donors (ages 81-84, 2 male, 2 female) were obtained from Proteogenex in a formalin fixed paraffin embedded (FFPE)-prepared form for IHC. Brain tissues were sectioned (7μm) and mounted onto super adhesive slides, dewaxed, and rehydrated following standard histology methods. Brain sections were then incubated with Tris-EDTA pH9 for 20 minutes in a programmable pressure cooker (Instant Pot™) for proper epitope retrieval. Following epitope retrieval, brain sections were photobleached overnight by exposing sections to a broad-spectrum LED light (ex/Aibecy A4 Ultra Bright 25,000 Lux LED Light Box-Tracing Pad) while being immersed in a solution of 4.5% (w/v) $H_2O_2$ and 20mM NaOH in PBS (*68*). The PhenoCycler Fusion platform (Akoya Biosciences, Marlborough, MA) was implemented for tissue analysis; brain sections were stained with a mixture of oligonucleotide barcoded PhenoCycler antibodies (CD31-RX001-488 (AKOYA, clone EP3950), α-SMA-RX013-488 (AKOYA, clone 1A4), CLDN-5-RX049-488 (Abcam, clone EPR7583), NeuN-RX019-488 (Invitrogen), TfR-RX046-550 (Abcam, clone EPR20584)), post-fixed according to the manufacturer's protocol, and imaged via the PhenoCycler Fusion system with a 20x long working distance objective (NA of 0.75) and a Sony IMX421-based camera.

**Microscopy image analysis**:

**2D mouse images.** Objects for imaging analysis (total tissue, total vessels, parenchyma, TfR, huIgG, and Aβ plaques) were identified using a custom Zeiss Zen image analysis script. 'Total tissue' was determined by standard thresholding in fluorescence channels and removing areas of folds or bubbles, leaving only significant areas of tissue with specific size restrictions. All individual fluorescence channel objects (CD31, TfR, huIgG, and Aβ plaques) were identified based on standard and dynamic (local) threshold levels and were only selected if identified within the defined 'total tissue' object. 'Total vessel' object was then identified by combining huIgG and CD31 objects together with objects exhibiting vessel-like morphology (using custom morphology software tools). 'Parenchyma' areas were defined as tissue areas not covered by either 'total vessel' or NeuN objects. Due to high overall staining intensity of both the TfR and huIgG channels (AF647) in neonates, image thresholds were adjusted to allow similar object detection across all ages: high/more stringent thresholds in neonates, and lower – less stringent thresholds in juveniles and adults. All other object settings (total tissue, total vessels, parenchyma, and individual channel objects) were held constant across all tissues analyzed. Lastly, Zen was also used to quantify total areas and total sum signal intensities of various fluorescence channels across each class of identified objects. All Zeiss Axioscan.Z1 images with the respective identified objects were loaded into napari v0.5.2 (*70*) for visualization and region of interest (ROI) selection. Cortical ROIs were selected directly superior to the hippocampus for all animals. Stark differences in signal intensity for TfR and huIgG in neonates required separate contrast limits than those of juveniles and adults. To enable qualitative comparison across age, a one parameter gamma distribution was fit for the histograms in all images using scipy v1.13.0 (*71*) and averaged per age, giving an ~4x increase of the scale parameter in the neonates compared to juveniles and adults. Similarly, image generation for the neonatal brain required different visualization settings (the right window of the contrast limit for juveniles and adults were scaled by 4.0) than those of the other age groups. Differences in TfR, huIgG, or Aβ were analyzed across age using one-way ANOVA, Tukey's post hoc in GraphPad Prism v10.3.1.

**3D mouse hemibrain images.** Lightsheet image TIFs were converted to an Imaris file format (IMS) and stitched together for visualization and analysis in Imaris v10.1.1 (Oxford Instruments). Using Imaris machine learning-aided surface segmentation, TfR signal was extracted from the vascular and perivascular space of each hemibrain. 'Total Vascular Volume' was determined by first combining both TfR and Lectin channels and then training the machine learning algorithm to create vascular/perivascular segmentation by providing repeated examples of foreground vascular space and background non-vascular space from animals representing each adult age group (3-, 15-, and 21.5-month-old mice). The final vascular/perivascular segmented surface was filtered to exclude all small (less than 10μm diameter objects) or round (objects with greater than 0.7 sphericity), leaving longer vascular objects within the surface. Following segmentation, absolute TfR signal was assessed across ages within the vascular/perivascular surface, where the mean TfR intensity was determined by dividing the total TfR intensity within the surface by the vascular/perivascular surface volume was analyzed by one-way ANOVA, Tukey's post hoc in GraphPad Prism v10.3.1.

**Human CODEX images.** All raw fluorescent channels of a given CODEX image were converted to a zarr image pyramid (*72*) by down sampling each image in ITK (*73*). Unless specified otherwise, the final full resolution image was used to generate segmentation masks and quantifications. To begin, a 'whole-tissue' mask was first defined by thresholding DAPI followed by the removal of holes and segmented objects smaller than 500 pixel$^2$ (~4,100 μm$^2$). Only values within the defined whole-tissue mask were considered for any further segmentation of substructures or quantification of image intensities. Gray and white matter substructures were next defined by first calculating a foreground NeuN image (i.e., subtracting the background of the original NeuN image from a blurred version of the full resolution NeuN image) and classifying pixels above the individual thresholds of both the raw and foreground NeuN images, as 'neurons'. Segmented neurons smaller than 100 pixels$^2$ (~26 μm$^2$) were then filtered from the 'neuron' mask, and a 1 pixel (~4 μm) binary closing was performed to merge the 'neuron' mask to the gray matter. In the 'merged neuron' mask, holes smaller than 5 pixels$^2$ (~10 μm$^2$) were filled and objects smaller than 50 pixels$^2$ (~100 μm$^2$) were removed, leaving the resulting pixels to be defined as 'gray matter'. All other pixels within the 'whole-tissue mask' that did not merge with the 'gray matter' mask, were defined as 'white matter'. Each 'gray matter' or 'white matter' mask was then manually inspected at full resolution and the segmentation boundaries were manually corrected to remove sampling and edge artifacts with the Label tools in napari (*70*). Next, a 'vessel' mask was generated by thresholding both the original and foreground images for each vascular marker (α−SMA, CD31, and Claudin-5), accepting only pixels over both thresholds, then forming the new vessel mask as the union of all three vascular channels. Vessels were then split into contiguous objects using scipy.ndimage.label (*71*), and α−SMA$^+$ vessels (i.e., arteries) were classified as any vessel where >10% of the individual object mask overlapped with the α−SMA$^+$ mask. All remaining vessels were classified as α−SMA$^-$ (i.e., veins, venules, and capillaries), and any pixels in the remaining 'whole-tissue' mask were defined as 'parenchyma'. To refine vessel boundaries and reduce sensitivity to background inhomogeneity, the vessel segmentation was further refined with a random forest classifier (scikit-learn v1.4.0) (*74*) to predict a probability mask for each α−SMA$^+$ vessel, α−SMA$^-$ vessel, or parenchymal object. The probability masks for α−SMA$^+$ and α−SMA$^-$ vessels were thresholded at 10% and 15% probability respectively, then the union of the two were used as the final vessel mask. Vessels were again split into contiguous objects using scipy.ndimage.label, then α−SMA$^+$ vessels were classified as any vessel where >10% of the individual object mask overlapped with the α−SMA$^+$ mask. The remaining unclassified pixels were defined as parenchyma, regardless of the label assigned by the random forest classifier. Lastly, the mean intensity for the individual TfR channel was compared between AD and control brains across α−SMA$^+$ vessels, α−SMA$^-$ vessels, and the brain parenchyma in both gray and white compartments using a one-way ANOVA, Tukey's post hoc test, where significant differences were assessed at $p<0.05$ using two-way ANOVA, Tukey's post hoc in GraphPad Prism v10.3.1.

**Capillary depletion:** As previously described (*57*), fresh, PBS-perfused brains were grossly dissected, and both the meninges and choroid plexus were removed via additional microdissection. Using a dounce homogenizer, the microdissected brains were then homogenized in HBSS followed by centrifugation at 1,000G for 10 minutes at 4°C. The 'non-cell associated fraction' supernatant was removed (and further centrifuged at 14,000G for 10 minutes at 4°C to isolate a clean non-cell associated fraction) and cell pellets were resuspended in 17% dextran for gradient centrifugation to separate brain vasculature from all other parenchymal cells. An aliquot of the resuspended cells was collected, washed in HBSS, lysed for 30 seconds at 27Hz (Qiagen TissueLyser II) at 4°C in lysis buffer (1% NP40-PBS containing protease and phosphatase inhibitors (Roche)), and further centrifuged at 14,000G for 10 minutes at 4°C to isolate the total cell-associated lysate. The remaining cells resuspended in 17% dextran proceeded to centrifugation at 4,122G for 15 minutes at 4°C. The 17% dextran-induced gradient centrifugation yielded a supernatant containing myelin and parenchymal cells and a cell pellet containing vascular cells. The myelin + parenchymal cell supernatant was collected, washed with HBSS, and centrifuged at 4,122G for 15 minutes at 4°C, yielding a parenchymal cell pellet. Both vascular and parenchymal cell pellets were them subsequently lysed in 150 or 500µL, respectively, further centrifuged at 14,000G for 10 minutes at 4°C, and the protein concentrations of the resulting lysates were determined by BCA (Thermo A55860) according to the manufacturer's protocol. All samples were then stored at -80°C until further analysis. Any IgG measurements within the vascular and parenchymal fractions are presented as 'absolute concentrations' as we observed age-related changes in total protein concentrations in these fractions that would confound sample analyses.

**TfR MSD ELISA:** TfR MSD (Meso Scale Discovery) was performed as previously described (*34*). Briefly, using automated robotics technology and constant plate agitation at 700RPM, 384-well MSD Gold Streptavidin plates were blocked for 1 hour at room temperature (RT) with 5% MSD blocker, washed with PBST, coated for 1 hour at RT with 0.125µg/mL of an in-house biotinylated TfR capture antibody (part of Abcam kit ab25663), and subsequently incubated with a working concentration of sample (1:300 for brain lysate and 1:2 for BEC cell lysate) overnight at 4°C. Plates were then washed, incubated for 1 hour at RT with 0.25µg/mL of an in-house sulfo-tagged TfR detection antibody (part of Abcam kit ab25663), washed again, and incubated for 10-15 minutes at RT with 2x MSD read buffer and read on an MSD plate reader (Meso Sector S 600) to acquire data. MSD data were analyzed one- or two-way ANOVA, Tukey's hoc in GraphPad Prism v10.3.1 and where specified, mouse TfR concentrations were either normalized to the total protein concentration (determined by BCA assay (Thermo A55860) according to the manufacturer's protocol) or the cell number (determined by FACS) of each individual sample.

**Antibody and ATV$^{TfR}$ generation:** As previously described (*14*), the light and heavy chain sequences of control huIgG or the ATV$^{TfR}$ variants were cloned into a pRK5 expression vector, using anti-DNP02 Fabs, and expressed in Expi293F™ cells (Gibco/Thermo Fisher A14527 using standard methods. Using protein A chromatography (Mab Select SuRe, Cytiva), both control huIgG and ATV$^{TfR}$ variants were first affinity purified from clarified culture supernatants, followed by cation exchange chromatography (CaptoS Impact, Cytiva), and then stored in at 4°C in PBS or standard formulation buffer (10mM sodium acetate with 6% sucrose, pH5.5). The identities of purified, intact control huIgG or ATV$^{TfR}$ molecules were confirmed by LC/MS, and a purity of >95% was confirmed by CE-SDS and analytical HPLC-SEC.

**IgG quantification and analysis:**

Plasma, CSF and brain muIgG and huIgG concentrations were quantified using a generic anti-mouse IgG sandwich-format ELISA or anti-human IgG sandwich-format ELISA (*9*), respectively. Due to low concentrations of huIgG in the therapeutically relevant $ATV^{TfR}$ dose studies (10mg/kg in Figure 3 and Supplemental Figure 2), a generic huIgG MSD-based ELISA assay with increased huIgG sensitivity was used for those samples. All standard ELISA and MSD ELISA reaction steps were performed at RT unless otherwise specified, with gentle agitation where appropriate.

**muIgG ELISA.** Plates were coated overnight at 4°C with 0.5μg/mL donkey anti-mouse IgG (JIR #715-005-150) diluted in sodium bicarbonate solution (Sigma #C3041-50CAP), with no agitation. Following the overnight incubation, plates were washed 4x (PBS + 0.05% Tween 20) and incubated for 2 hours at RT with block buffer (PBS + 0.05% Tween 20 + 5% BSA). Plates were then washed 4x and incubated for 2 hours at RT with diluted assay standards or samples. Plasma, CSF, and brain samples were diluted 1:20,000, 1:40, and 1:10 in dilution buffer (PBS + 0.05% Tween 20 and 1% BSA), respectively. After sample incubation, plates were washed 4x and 0.02μg/mL of the detection goat anti-mouse IgG antibody (JIR #115-035-071) diluted in dilution buffer and was incubated for 1 hour at RT. Plates were again washed 4x and then developed for 5-10 minutes with TMB substrate (Thermo # 34029). The reaction was quenched with 2N sulfuric acid ($H_2SO_4$) and plates were read at 450 nm absorbance. Lastly, muIgG concentrations were calculated from muIgG ELISA standard curve (on average, the range for plasma and CSF was 0.006-3.03mg/mL and for brain was 0.328-330μg/mL) and analyzed by one- or two-way ANOVA, Tukey's hoc in GraphPad Prism v10.3.1.

**huIgG ELISA.** As previously described (*9*), plates were coated overnight at 4°C with 1μg/mL donkey anti-human IgG (JIR #709-006-098) diluted in sodium bicarbonate solution (Sigma #C3041-50CAP), washed 3x, and blocked for 2 hours at RT. Plates were then washed 3x and incubated for 2 hours at RT with diluted assay standards or samples. Plasma, CSF, and brain samples were diluted 1:20,000, 1:10, and 1:20, respectively, in dilution buffer (PBS + 0.05% Tween 20 and 1% BSA). After incubation, plates were washed 3x and 0.02μg/mL of the detection goat anti-human IgG antibody (JIR #109-036-098) diluted in dilution buffer and was incubated for 1 hour at RT. Plates were again washed 3x and then developed for 5-10 minutes with TMB substrate. The reaction was quenched with 4N sulfuric acid ($H_2SO_4$) and plates were read at 450 nm absorbance. Lastly, huIgG concentrations were calculated from the huIgG ELISA standard curve (ranged from 0.001-300ng/mL) and analyzed by one- or two-way ANOVA, Tukey's hoc in GraphPad Prism v10.3.1.

**huIgG MSD ELISA.** MSD GOLD 96-well small-spot streptavidin-coated microtiter plates (Meso Scale Discovery, Rockville, MD) were blocked with 1% casein-based PBS (Thermo Scientific, Waltham, MA) for 1 hour at RT. The assay plates were then washed and incubated with a 0.5μg/mL working concentration solution of capture biotinylated goat anti-human IgG polyclonal antibody (SouthernBiotech, Birmingham, AL) in 1% casein-based PBS for 1-2 hours at RT. After a wash step, pre-diluted standards, quality controls, and study samples (including plasma, CSF, or brain lysate) were added to the assay plates and incubated for 1-2 hours at RT. During the sample incubation, a ruthenylated (SULFO-TAG) goat anti-human IgG polyclonal detection antibody (Meso Scale Discovery, Rockville, MD) was pre-adsorbed with blank matrices corresponding to the study samples being analyzed for 1 hour prior to its use in the assay plates. After the sample incubation followed by a wash step was completed, the plates were incubated with a 0.5µg/mL working concentration of the pre-adsorbed detection ruthenylated goat anti-human IgG antibody in 1% casein-based PBS for another 1 hour at RT. Following a final wash step, a 1x MSD Read Buffer T (Meso Scale Discovery, Rockville, MD) was added to the assay plates and were read on an MSD plate reader (Meso Sector S 600) to acquire data. The plasma and CSF assays had a standard curve with a dynamic range of 4.88-5000 ng/mL and a minimum-required-dilution (MRD) of 100. The brain lysate assay has a standard dynamic range of 2.44-2500 ng/mL and an MRD of 50. The raw data was processed into sample concentrations based on the assay standard curve fitted with a weighed four-parameter non-linear logistic regression using MSD Discovery Workbench 4.0 (Meso Scale Discovery, Rockville, MD) and analyzed by one- or two-way ANOVA, Tukey's hoc in GraphPad Prism v10.3.1.

**PK Parameters**. Non-compartmental analysis was estimated using Dotmatics software 5.5 (Boston, Mass) using intravenous route of administration with a linear up log down method. Parameters were estimated using nominal sampling times relative to the start of each administration. Samples that were below the quantitation limit (BQL) were omitted. Descriptive statistics (mean and standard deviation) were generated using Dotmatics.

**Supplemental Figures:**

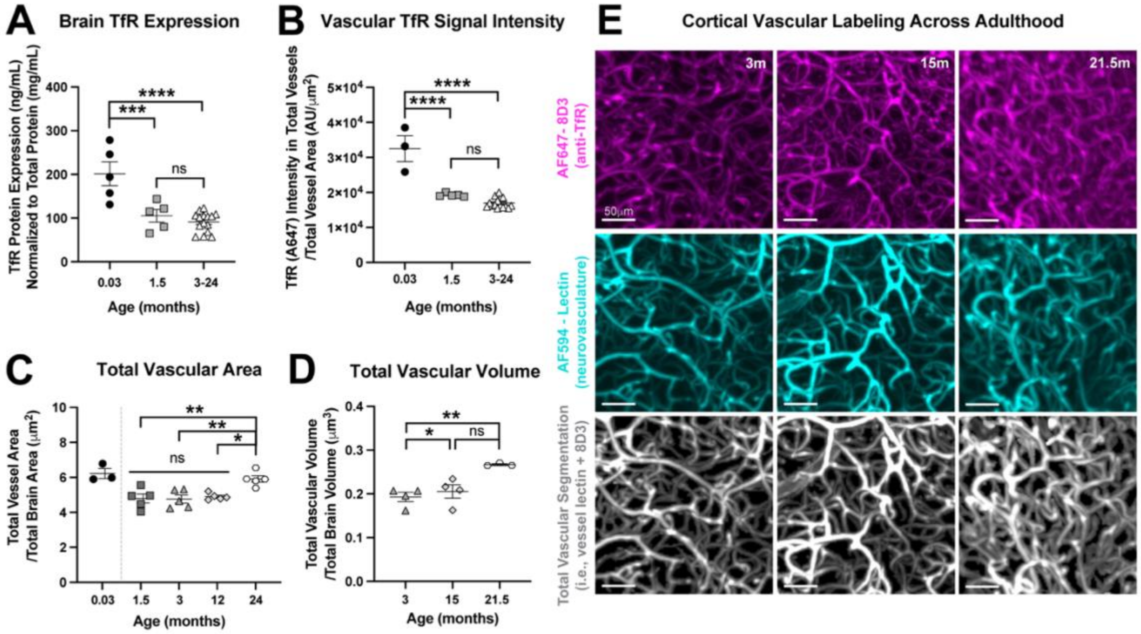


**Fig. S1. TfR is differentially expressed throughout healthy aging.** (**A**) Whole brain TfR protein expression quantified by MSD and normalized to total protein in bulk brain lysate of TfR$^{mu/hu}$ KI mice across 3 major developmental stages: neonate (0.03 months), juvenile (1.5 months), and adult (pooled average over 3-24 months). (**B**) Assessment of mean TfR intensity in sagittal brain sections of TfR$^{mu/hu}$ KI mice across segmented vasculature. Due to significantly high TfR expression and immunoreactivity in neonates, neonates were excluded from all statistical analyses in Fig. 1 to allow for the careful evaluation and detection of subtle changes in TfR levels within 1.5-24 month-old mice. (**C-D**) An age-related increase is observed in total vascular area of sagittal brain sections of TfR$^{mu/hu}$ KI mice (C) or total vascular volume in tissue-cleared hemibrains of WT mice (D). (**E**) Representative cortical images from tissue-cleared hemibrains of WT mice systemically dosed with AF647-labeled 8D3, targeted against murine TfR, and AF594-labeled lectin to label the neurovasculature and produce a 3D total vascular segmentation mask (i.e., merged 8D3 (AF647) and lectin (AF594) channels; scale bar 50μm)). (**A-D**) one-way ANOVA, Tukey's post hoc, graphs display mean ± SEM, n=3-5. * $p<0.05$, ** $p<0.01$, *** $p<0.001$, **** $p<0.0001$, see Tables S1-S2 for exact p values of all group comparisons.

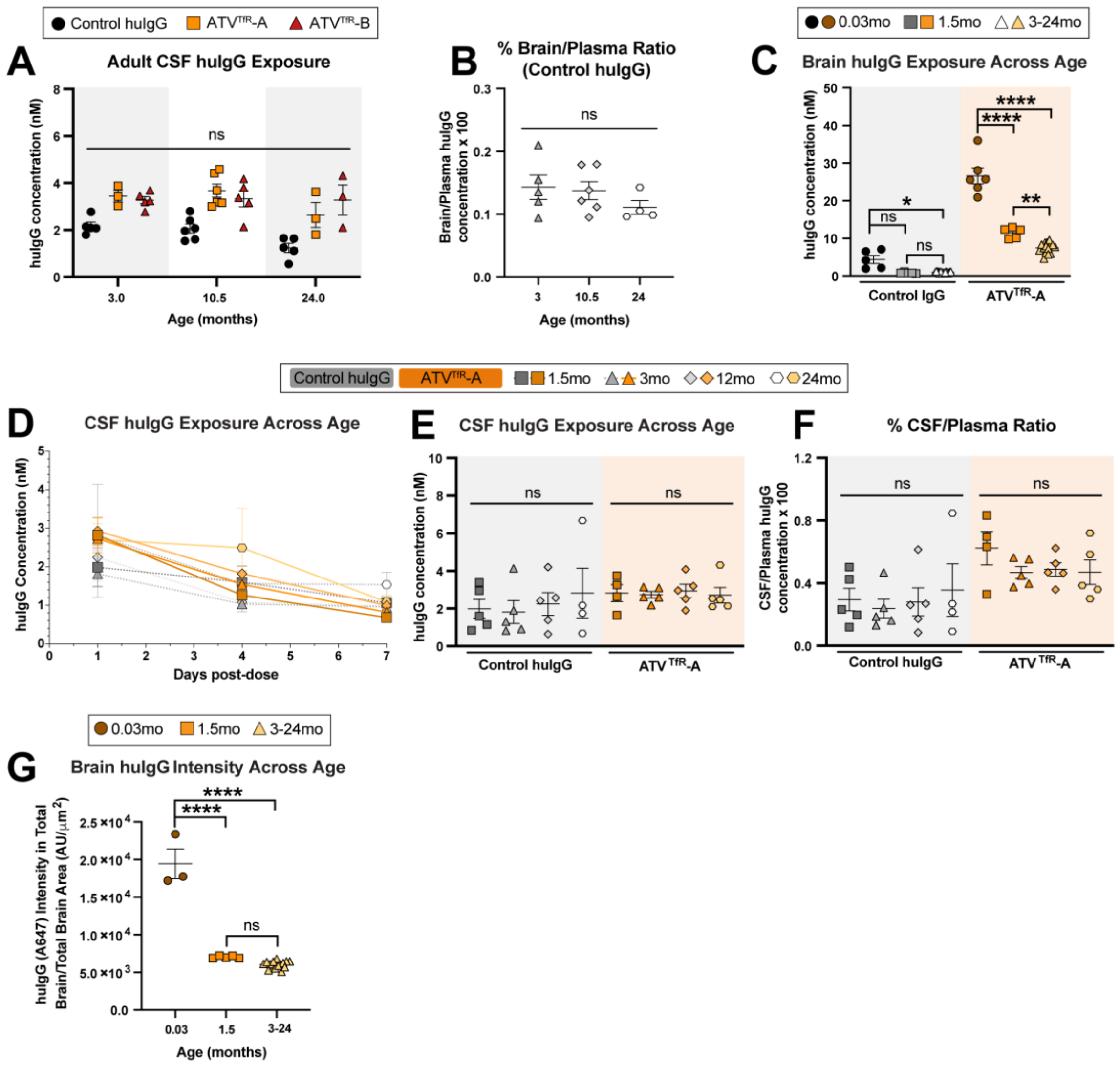


**Fig. S2**. **Neither CSF nor brain huIgG exposure are impacted across adult aging.** (**A**) absolute huIgG exposure in the CSF of TfR$^{mu/hu}$ KI mice 24 hours after a single 25 mg/kg i.v. dose of control huIgG, ATV$^{TfR}$-A, or ATV$^{TfR}$-B. (**B**) Brain-to-plasma ratio of the control huIgG group shown in Fig. 2c. (**C**) absolute huIgG concentrations in bulk brain lysate 24 hours after a single 10 mg/kg i.v. dose of either control huIgG or ATV$^{TfR}$-A across 3 major developmental stages: neonates (0.03 months), juveniles (1.5 months), and adults (pooled average over 3-24 months). The significantly high neonatal huIgG exposure in (C) was excluded from all statistical analyses in Fig. 3 to allow for the careful evaluation and detection of subtle changes in huIgG exposure within 1.5-24 month-old mice. (**D**) absolute huIgG concentrations over 7 days of ATV$^{TfR}$-A in the CSF of TfR$^{mu/hu}$ KI mice. (**E-F**) absolute huIgG CSF exposure (E) and CSF-to-plasma ratio (F) 24 hours after a single 10 mg/kg i.v. dose in TfR$^{mu/hu}$ KI mice. (**G**) Mean huIgG intensity quantified in sagittal brain sections of neonates, juveniles, and adults (pooled average over 3-24 months). The significantly high neonatal huIgG intensity in (G) was similarly excluded from all statistical analyses in Fig. 3 to allow for the careful evaluation and detection of subtle changes in huIgG intensity within 1.5-24 month-old mice. (**A, C, E-G**) two-way ANOVA, (**B, G**) one-way ANOVA, (**D**) see mean results in Table S1. (**A-G**) Graphs display mean ± SEM, n=4-6. * $p<0.05$, ** $p<0.01$, **** $p<0.0001$, see Tables S1 and S3 for exact p values of all group comparisons. $ATV^{TfR}$-A hTfR = ~1100nM $K_D$; $ATV^{TfR}$-B = hTfR ~100nM $K_D$

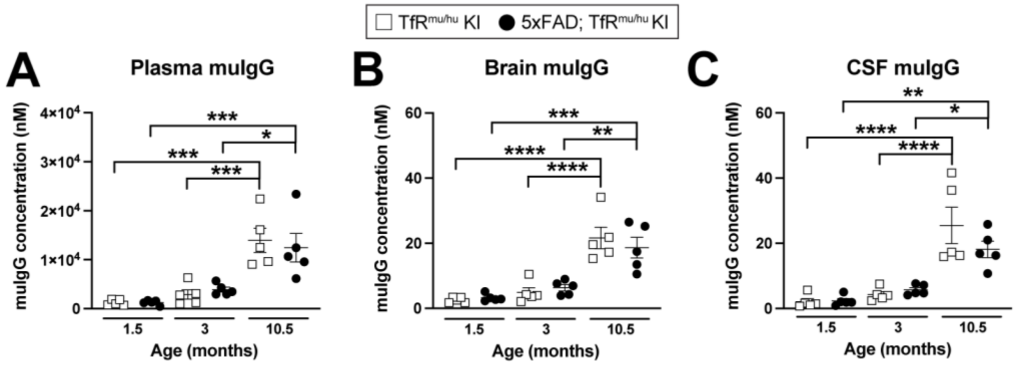


**Fig. S3. Disease progression does not impact endogenous muIgG levels in 5xFAD;TfR$^{mu/hu}$ KI mice.** (**A-C**) Quantification of endogenous muIgG levels in plasma (A), brain (B) and CSF (C) in 5xFAD;TfR$^{mu/hu}$ KI mice. (**A-C**) Two-way ANOVA, Tukey's post hoc, where all graphs display mean ± SEM, n=3-5; * $p<0.05$, ** $p<0.01$, *** $p<0.001$, **** $p<0.0001$, with only relevant comparisons indicated on graph; see Table S4 for exact p values of all group comparisons.

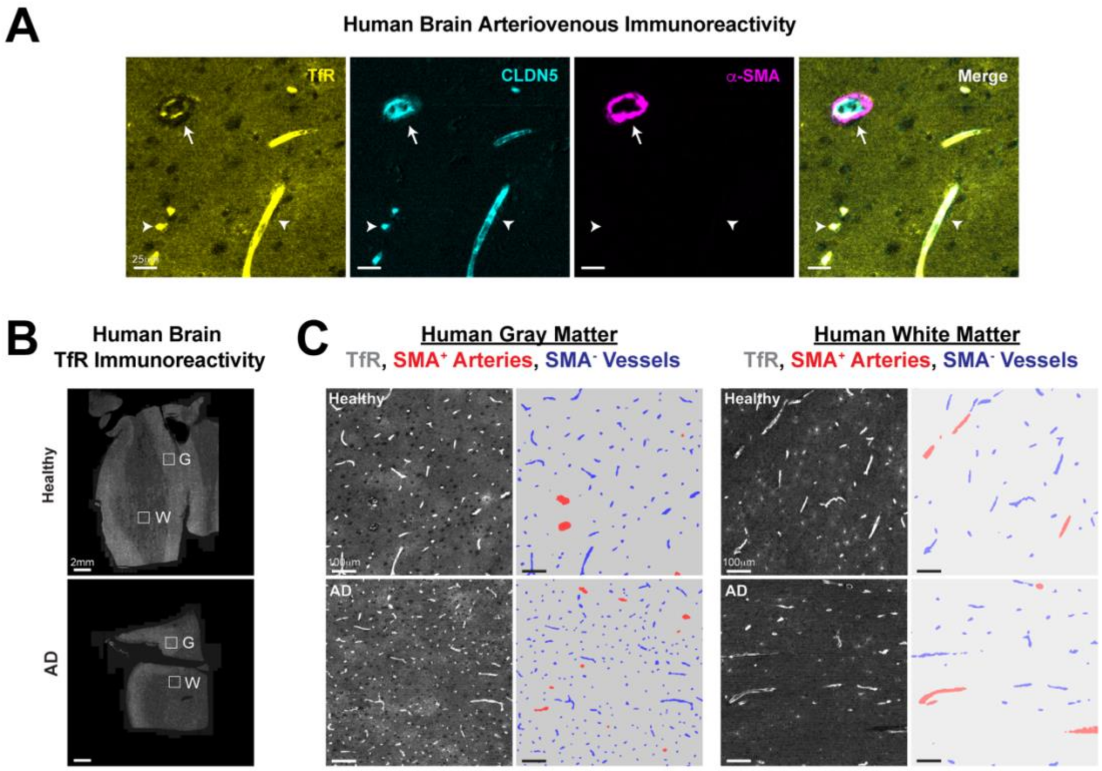


**Fig. S4**. **Alzheimer's disease does not impact TfR expression in human brain vasculature or parenchyma.** (**A**) Representative image from the grey matter of the frontal cortex in an 84-year-old healthy aged human brain qualitatively illustrating the expected zonal pattern of TfR protein along the arteriovenous axis (TfR, CLDN5, and α-SMA; scale bar 25μm). Arrow identifies large caliber (≥20μm) arteries/arterioles, whereas arrowheads identify smaller caliber microvessels (<10μm). (**B**) Representative images of TfR expression in healthy aged (top; 84 years old) and AD (bottom; 87 years old) human brain tissue quantified by CODEX (see Supplemental Materials and Methods; grey and white matter areas indicated by white squares; scale bar, 2mm). (**C**) Zoomed insets of the indicated grey and white matter areas (left; scale bar 100μm) and the respective automated vascular segmentation (right). Representative images are shown from n=4 humans/group, n=1 brain tissue sample per human; see Table S5 for human patient demographics.

### Supplemental Tables

**Table S1.** Overall statistical comparisons for all figures. Two-way ANOVA statistics are of interaction between age and treatment. Values below the lower limit of quantification are indicated as 'BLLOQ'.

| | **ANOVAs** | | | | |
|---|---|---|---|---|---|
| **Figure and Panel** | **Type** | **Sample or Segmentation Type** | **F** | **DF** | **P value** |
| Figure 1a | One-way | Bulk brain lysate | 0.96 | 3 | 0.4354 |
| Figure 1c | One-way | Vascular segmentation | 15.42 | 3 | 5.58E-05 |
| Figure 1d | One-way | Brain endothelial cell lysate | 15.57 | 3 | 5.26E-05 |
| Figure 1e | One-way | Vascular segmentation | 4.158 | 2 | 5.78E-02 |
| Figure 2a | Two-way | Plasma | 1.469 | 4 | 0.2307 |
| Figure 2b | Two-way | Bulk brain lysate | 1.984 | 4 | 0.1172 |
| Figure 2c | Two-way | % Bulk brain lysate:Plasma | 0.3971 | 4 | 0.8094 |
| Figure 2d | Two-way | % CSF:Plasma | 0.5729 | 4 | 0.6842 |
| Figure 3c | Two-way | % Bulk brain lysate:Plasma | 30.99 | 3 | 1.38E-09 |
| Figure 3f | Two-way | Non-cell-associated segmentation | 1.169 | 3 | 0.3389 |
| Figure 3h | One-way | Brain segmentation | 10.07 | 3 | 5.74E-04 |
| Figure 3i | One-way | Vascular segmentation | 7.137 | 3 | 2.94E-03 |
| Figure 3j | One-way | Parenchymal segmentation | 9.944 | 3 | 6.11E-04 |
| Figure 4a | One-way | % Ab plaque:Brain segmentation | 14.42 | 2 | 5.11E-03 |
| Figure 4b | Two-way | Bulk brain lysate | 0.9584 | 2 | 0.3976 |
| Figure 4c | Two-way | Brain endothelial cell lysate | 8.778 | 2 | 2.18E-03 |
| Figure 4d | Two-way | Plasma | 1.386 | 4 | 0.2625 |
| Figure 4e | Two-way | Bulk brain lysate | 2.181 | 4 | 9.52E-02 |
| Figure 4f | Two-way | Vascular lysate | 0.7688 | 2 | 0.4746 |
| Figure 4g | Two-way | Parenchymal lysate | 0.5693 | 2 | 0.5733 |
| Figure 4h | Two-way | % Bulk brain lysate:Plasma | 5.884 | 4 | 1.28E-03 |
| Figure 4i | Two-way | % CSF:Plasma | 1.332 | 4 | 0.2808 |
| Figure 4j | Two-way | % Bulk brain lysate:Plasma | 0.8107 | 2 | 0.4563 |
| Figure 4k | Two-way | % CSF:Plasma | 0.5914 | 2 | 0.5614 |
| Figure 4l | One-way | Human vasc. & parench. segmentation | 26.39 | 11 | 5.70E-14 |
| Supp. Figure 1a | One-way | Bulk brain lysate | 19.59 | 2 | 1.31E-05 |
| Supp. Figure 1b | One-way | Vascular segmentation | 54.83 | 2 | 7.63E-09 |
| Supp. Figure 1c | One-way | Vascular segmentation | 7.487 | 3 | 2.38E-03 |
| Supp. Figure 1d | One-way | Vascular segmentation | 9.832 | 2 | 6.99E-03 |
| Supp. Figure 2a | Two-way | CSF | 0.9322 | 4 | 0.4577 |

| | | | | | |
|---|---|---|---|---|---|
| Supp. Figure 2b | One-way | % Bulk brain lysate:Plasma | 1.025 | 2 | 0.3883 |
| Supp. Figure 2c | Two-way | Bulk brain lysate | 58.17 | 2 | 3.34E-13 |
| Supp. Figure 2e | Two-way | CSF | 0.2992 | 3 | 0.8256 |
| Supp. Figure 2f | Two-way | % CSF:Plasma | 0.5137 | 3 | 0.6759 |
| Supp. Figure 2g | One-way | Brain segmentation | 170.8 | 2 | 2.69E-13 |
| Supp. Figure 3a | Two-way | CSF | 0.8400 | 4 | 0.5107 |
| Supp. Figure 3b | Two-way | Plasma | 0.2940 | 2 | 0.7479 |
| Supp. Figure 3c | Two-way | Bulk brain lysate | 0.6956 | 2 | 0.5085 |
| Supp. Figure 3d | Two-way | CSF | 1.7702 | 2 | 0.2035 |
| **Mean Results (Time course studies)** | | | | | |
| **Figure and Panel** | **Treatment and Age** | **Sample type and Day post-dose** | **Mean and Unit** | **SD** | **Group size 'n'** |
| Figure 3a | Control huIgG 1.5 months | Plasma 1-day post-dose | 0.6688 mM | 0.0519 | 5 |
| Figure 3a | Control huIgG 3 months | Plasma 1-day post-dose | 0.7115 mM | 0.1267 | 5 |
| Figure 3a | Control huIgG 12 months | Plasma 1-day post-dose | 0.8266 mM | 0.1178 | 5 |
| Figure 3a | Control huIgG 24 months | Plasma 1-day post-dose | 0.8083 mM | 0.0558 | 5 |
| Figure 3a | $ATV^{TfR}$-A 1.5 months | Plasma 1-day post-dose | 0.4600 mM | 0.0304 | 5 |
| Figure 3a | $ATV^{TfR}$-A 3 months | Plasma 1-day post-dose | 0.5871 mM | 0.0398 | 5 |
| Figure 3a | $ATV^{TfR}$-A 12 months | Plasma 1-day post-dose | 0.6083 mM | 0.1574 | 5 |
| Figure 3a | $ATV^{TfR}$-A 24 months | Plasma 1-day post-dose | 0.5938 mM | 0.1021 | 5 |
| Figure 3a | Control huIgG 1.5 months | Plasma 4-days post-dose | 0.5571 mM | 0.0308 | 5 |
| Figure 3a | Control huIgG 3 months | Plasma 4-days post-dose | 0.6598 mM | 0.0269 | 5 |
| Figure 3a | Control huIgG 12 months | Plasma 4-days post-dose | 0.6852 mM | 0.0373 | 5 |
| Figure 3a | Control huIgG 24 months | Plasma 4-days post-dose | 0.6991 mM | 0.1734 | 5 |
| Figure 3a | $ATV^{TfR}$-A 1.5 months | Plasma 4-days post-dose | 0.1903 mM | 0.0099 | 5 |
| Figure 3a | $ATV^{TfR}$-A 3 months | Plasma 4-days post-dose | 0.2785 mM | 0.0319 | 5 |
| Figure 3a | $ATV^{TfR}$-A 12 months | Plasma 4-days post-dose | 0.3054 mM | 0.0335 | 5 |
| Figure 3a | $ATV^{TfR}$-A 24 months | Plasma 4-days post-dose | 0.2781 mM | 0.0636 | 5 |
| Figure 3a | Control huIgG 1.5 months | Plasma 7-days post-dose | 0.4609 mM | 0.0364 | 5 |
| Figure 3a | Control huIgG 3 months | Plasma 7-days post-dose | 0.5432 mM | 0.0798 | 5 |
| Figure 3a | Control huIgG 12 months | Plasma 7-days post-dose | 0.5386 mM | 0.1970 | 5 |
| Figure 3a | Control huIgG 24 months | Plasma 7-days post-dose | 0.5493 mM | 0.0441 | 5 |
| Figure 3a | $ATV^{TfR}$-A 1.5 months | Plasma 7-days post-dose | 0.0845 mM | 0.0158 | 5 |
| Figure 3a | $ATV^{TfR}$-A 3 months | Plasma 7-days post-dose | 0.1375 mM | 0.0109 | 5 |
| Figure 3a | $ATV^{TfR}$-A 12 months | Plasma 7-days post-dose | 0.1475 mM | 0.0440 | 5 |
| Figure 3a | $ATV^{TfR}$-A 24 months | Plasma 7-days post-dose | 0.1610 mM | 0.040 | 4 |

| | | | | | |
|---|---|---|---|---|---|
| Figure 3b | Control huIgG 0.03 months | Bulk brain lysate 1-day post-dose | 4.3882 nM | 2.3932 | 5 |
| Figure 3b | Control huIgG 1.5 months | Bulk brain lysate 1-day post-dose | 0.8656 nM | 0.1886 | 5 |
| Figure 3b | Control huIgG 3 months | Bulk brain lysate 1-day post-dose | 0.9088 nM | 0.1938 | 5 |
| Figure 3b | Control huIgG 12 months | Bulk brain lysate 1-day post-dose | 0.9754 nM | 0.1666 | 5 |
| Figure 3b | Control huIgG 24 months | Bulk brain lysate 1-day post-dose | 1.1084 nM | 0.1071 | 5 |
| Figure 3b | $ATV^{TfR}$-A 0.03 months | Bulk brain lysate 1-day post-dose | 26.6261 nM | 5.1823 | 6 |
| Figure 3b | $ATV^{TfR}$-A 1.5 months | Bulk brain lysate 1-day post-dose | 11.501 nM | 1.4190 | 5 |
| Figure 3b | $ATV^{TfR}$-A 3 months | Bulk brain lysate 1-day post-dose | 8.69 nM | 0.7177 | 5 |
| Figure 3b | $ATV^{TfR}$-A 12 months | Bulk brain lysate 1-day post-dose | 6.6922 nM | 1.3632 | 5 |
| Figure 3b | $ATV^{TfR}$-A 24 months | Bulk brain lysate 1-day post-dose | 7.0694 nM | 1.1608 | 5 |
| Figure 3b | Control huIgG 0.03 months | Bulk brain lysate 4-days post-dose | 1.9968 nM | 0.6911 | 5 |
| Figure 3b | Control huIgG 1.5 months | Bulk brain lysate 4-days post-dose | 0.7466 nM | 0.08789 | 5 |
| Figure 3b | Control huIgG 3 months | Bulk brain lysate 4-days post-dose | 0.6874 nM | 0.0656 | 5 |
| Figure 3b | Control huIgG 12 months | Bulk brain lysate 4-days post-dose | 0.92 nM | 0.1056 | 5 |
| Figure 3b | Control huIgG 24 months | Bulk brain lysate 4-days post-dose | 1.0402 nM | 0.4616 | 5 |
| Figure 3b | $ATV^{TfR}$-A 0.03 months | Bulk brain lysate 4-days post-dose | 8.967 nM | 0.6898 | 6 |
| Figure 3b | $ATV^{TfR}$-A 1.5 months | Bulk brain lysate 4-days post-dose | 5.4046 nM | 0.2083 | 5 |
| Figure 3b | $ATV^{TfR}$-A 3 months | Bulk brain lysate 4-days post-dose | 4.9446 nM | 0.5067 | 5 |
| Figure 3b | $ATV^{TfR}$-A 12 months | Bulk brain lysate 4-days post-dose | 4.9604 nM | 0.6937 | 5 |
| Figure 3b | $ATV^{TfR}$-A 24 months | Bulk brain lysate 4-days post-dose | 4.7784 nM | 0.7691 | 5 |
| Figure 3b | Control huIgG 0.03 months | Bulk brain lysate 7-days post-dose | 0.9102 nM | 0.2673 | 5 |
| Figure 3b | Control huIgG 1.5 months | Bulk brain lysate 7-days post-dose | 0.6336 nM | 0.1013 | 5 |
| Figure 3b | Control huIgG 3 months | Bulk brain lysate 7-days post-dose | 0.7208 nM | 0.1135 | 5 |
| Figure 3b | Control huIgG 12 months | Bulk brain lysate 7-days post-dose | 0.7346 nM | 0.2263 | 5 |
| Figure 3b | Control huIgG 24 months | Bulk brain lysate 7-days post-dose | 1.032 nM | 0.4442 | 5 |
| Figure 3b | $ATV^{TfR}$-A 0.03 months | Bulk brain lysate 7-days post-dose | 1.8188 nM | 0.2124 | 6 |
| Figure 3b | $ATV^{TfR}$-A 1.5 months | Bulk brain lysate 7-days post-dose | 2.8048 nM | 0.2820 | 5 |
| Figure 3b | $ATV^{TfR}$-A 3 months | Bulk brain lysate 7-days post-dose | 2.9624 nM | 0.2093 | 5 |
| Figure 3b | $ATV^{TfR}$-A 12 months | Bulk brain lysate 7-days post-dose | 2.8784 nM | 0.7197 | 5 |
| Figure 3b | $ATV^{TfR}$-A 24 months | Bulk brain lysate 7-days post-dose | 3.0295 nM | 0.4623 | 4 |
| Figure 3d | Control huIgG 0.03 months | Vascular lysate 1-day post-dose | 0.0630 nM | 0.0787 | 4 |
| Figure 3d | Control huIgG 1.5 months | Vascular lysate 1-day post-dose | 0.0273 nM | 0.0025 | 5 |
| Figure 3d | Control huIgG 3 months | Vascular lysate 1-day post-dose | 0.0278 nM | 0.0121 | 4 |
| Figure 3d | Control huIgG 12 months | Vascular lysate 1-day post-dose | 0.0308 nM | 0.0060 | 5 |
| Figure 3d | Control huIgG 24 months | Vascular lysate 1-day post-dose | 0.0251 nM | 0.0051 | 3 |

| | | | | | |
|---|---|---|---|---|---|
| Figure 3d | ATV$^{TfR}$-A 0.03 months | Vascular lysate 1-day post-dose | 1.2020 nM | 0.3425 | 6 |
| Figure 3d | ATV$^{TfR}$-A 1.5 months | Vascular lysate 1-day post-dose | 0.6159 nM | 0.1010 | 5 |
| Figure 3d | ATV$^{TfR}$-A 3 months | Vascular lysate 1-day post-dose | 0.4959 nM | 0.0590 | 5 |
| Figure 3d | ATV$^{TfR}$-A 12 months | Vascular lysate 1-day post-dose | 0.4222 nM | 0.1290 | 5 |
| Figure 3d | ATV$^{TfR}$-A 24 months | Vascular lysate 1-day post-dose | 0.4214 nM | 0.2174 | 5 |
| Figure 3d | Control huIgG 0.03 months | Vascular lysate 4-days post-dose | BLLOQ | BLLOQ | - |
| Figure 3d | Control huIgG 1.5 months | Vascular lysate 4-days post-dose | 0.0390 nM | 0.0157 | 4 |
| Figure 3d | Control huIgG 3 months | Vascular lysate 4-days post-dose | 0.0199 nM | 0.0006 | 4 |
| Figure 3d | Control huIgG 12 months | Vascular lysate 4-days post-dose | 0.0418 nM | 0.0369 | 3 |
| Figure 3d | Control huIgG 24 months | Vascular lysate 4-days post-dose | 0.0423 nM | 0.0226 | 4 |
| Figure 3d | ATV$^{TfR}$-A 0.03 months | Vascular lysate 4-days post-dose | 0.3901 nM | 0.1303 | 6 |
| Figure 3d | ATV$^{TfR}$-A 1.5 months | Vascular lysate 4-days post-dose | 0.2657 nM | 0.0500 | 5 |
| Figure 3d | ATV$^{TfR}$-A 3 months | Vascular lysate 4-days post-dose | 0.2586 nM | 0.0333 | 5 |
| Figure 3d | ATV$^{TfR}$-A 12 months | Vascular lysate 4-days post-dose | 0.3890 nM | 0.2479 | 5 |
| Figure 3d | ATV$^{TfR}$-A 24 months | Vascular lysate 4-days post-dose | 0.3601 nM | 0.1200 | 5 |
| Figure 3d | Control huIgG 0.03 months | Vascular lysate 7-days post-dose | BLLOQ nM | BLLOQ | - |
| Figure 3d | Control huIgG 1.5 months | Vascular lysate 7-days post-dose | 0.0290 nM | 0.0044 | 2 |
| Figure 3d | Control huIgG 3 months | Vascular lysate 7-days post-dose | 0.0348 nM | 0.0217 | 4 |
| Figure 3d | Control huIgG 12 months | Vascular lysate 7-days post-dose | 0.0260 nM | 0.0055 | 3 |
| Figure 3d | Control huIgG 24 months | Vascular lysate 7-days post-dose | 0.0378 nM | 0.0061 | 2 |
| Figure 3d | ATV$^{TfR}$-A 0.03 months | Vascular lysate 7-days post-dose | 0.1223 nM | 0.0573 | 6 |
| Figure 3d | ATV$^{TfR}$-A 1.5 months | Vascular lysate 7-days post-dose | 0.1067 nM | 0.0791 | 5 |
| Figure 3d | ATV$^{TfR}$-A 3 months | Vascular lysate 7-days post-dose | 0.2795 nM | 0.2863 | 5 |
| Figure 3d | ATV$^{TfR}$-A 12 months | Vascular lysate 7-days post-dose | 0.1345 nM | 0.0899 | 5 |
| Figure 3d | ATV$^{TfR}$-A 24 months | Vascular lysate 7-days post-dose | 0.4954 nM | 0.2438 | 4 |
| Figure 3e | Control huIgG 0.03 months | Parenchymal lysate 1-day post-dose | BLLOQ | BLLOQ | - |
| Figure 3e | Control huIgG 1.5 months | Parenchymal lysate 1-day post-dose | BLLOQ | BLLOQ | - |
| Figure 3e | Control huIgG 3 months | Parenchymal lysate 1-day post-dose | BLLOQ | BLLOQ | - |
| Figure 3e | Control huIgG 12 months | Parenchymal lysate 1-day post-dose | BLLOQ | BLLOQ | - |
| Figure 3e | Control huIgG 24 months | Parenchymal lysate 1-day post-dose | BLLOQ | BLLOQ | - |
| Figure 3e | ATV$^{TfR}$-A 0.03 months | Parenchymal lysate 1-day post-dose | 0.1543 nM | 0.0972 | 6 |
| Figure 3e | ATV$^{TfR}$-A 1.5 months | Parenchymal lysate 1-day post-dose | 0.2761 nM | 0.0944 | 5 |
| Figure 3e | ATV$^{TfR}$-A 3 months | Parenchymal lysate 1-day post-dose | 0.1991 nM | 0.1991 | 5 |
| Figure 3e | ATV$^{TfR}$-A 12 months | Parenchymal lysate 1-day post-dose | 0.1650 nM | 0.0287 | 5 |
| Figure 3e | ATV$^{TfR}$-A 24 months | Parenchymal lysate 1-day post-dose | 0.1517 nM | 0.0465 | 4 |

| | | | | | |
|---|---|---|---|---|---|
| Figure 3e | Control huIgG 0.03 months | Parenchymal lysate 4-days post-dose | BLLOQ | BLLOQ | - |
| Figure 3e | Control huIgG 1.5 months | Parenchymal lysate 4-days post-dose | BLLOQ | BLLOQ | - |
| Figure 3e | Control huIgG 3 months | Parenchymal lysate 4-days post-dose | BLLOQ | BLLOQ | - |
| Figure 3e | Control huIgG 12 months | Parenchymal lysate 4-days post-dose | BLLOQ | BLLOQ | - |
| Figure 3e | Control huIgG 24 months | Parenchymal lysate 4-days post-dose | BLLOQ | BLLOQ | - |
| Figure 3e | $ATV^{TfR}$-A 0.03 months | Parenchymal lysate 4-days post-dose | 0.2423 nM | 0.1112 | 6 |
| Figure 3e | $ATV^{TfR}$-A 1.5 months | Parenchymal lysate 4-days post-dose | 0.1455 nM | 0.0302 | 5 |
| Figure 3e | $ATV^{TfR}$-A 3 months | Parenchymal lysate 4-days post-dose | 0.1075 nM | 0.0187 | 5 |
| Figure 3e | $ATV^{TfR}$-A 12 months | Parenchymal lysate 4-days post-dose | 0.1152 nM | 0.0349 | 5 |
| Figure 3e | $ATV^{TfR}$-A 24 months | Parenchymal lysate 4-days post-dose | 0.1150 nM | 0.0306 | 5 |
| Figure 3e | Control huIgG 0.03 months | Parenchymal lysate 7-days post-dose | BLLOQ | BLLOQ | - |
| Figure 3e | Control huIgG 1.5 months | Parenchymal lysate 7-days post-dose | BLLOQ | BLLOQ | - |
| Figure 3e | Control huIgG 3 months | Parenchymal lysate 7-days post-dose | BLLOQ | BLLOQ | - |
| Figure 3e | Control huIgG 12 months | Parenchymal lysate 7-days post-dose | BLLOQ | BLLOQ | - |
| Figure 3e | Control huIgG 24 months | Parenchymal lysate 7-days post-dose | BLLOQ | BLLOQ | - |
| Figure 3e | $ATV^{TfR}$-A 0.03 months | Parenchymal lysate 7-days post-dose | 0.0326 nM | 0.0127 | 4 |
| Figure 3e | $ATV^{TfR}$-A 1.5 months | Parenchymal lysate 7-days post-dose | 0.0958 nM | 0.0198 | 5 |
| Figure 3e | $ATV^{TfR}$-A 3 months | Parenchymal lysate 7-days post-dose | 0.1246 nM | 0.0293 | 5 |
| Figure 3e | $ATV^{TfR}$-A 12 months | Parenchymal lysate 7-days post-dose | 0.0803 nM | 0.0199 | 5 |
| Figure 3e | $ATV^{TfR}$-A 24 months | Parenchymal lysate 7-days post-dose | 0.1158 nM | 0.0186 | 4 |
| Supp. Figure 2d | Control huIgG 1.5 months | CSF 1-day post-dose | 1.9864 nM | 1.1270 | 5 |
| Supp. Figure 2d | Control huIgG 3 months | CSF 1-day post-dose | 1.8164 nM | 1.3661 | 5 |
| Supp. Figure 2d | Control huIgG 12 months | CSF 1-day post-dose | 2.2447 nM | 1.3489 | 5 |
| Supp. Figure 2d | Control huIgG 24 months | CSF 1-day post-dose | 2.8189 nM | 2.6469 | 4 |
| Supp. Figure 2d | $ATV^{TfR}$-A 1.5 months | CSF 1-day post-dose | 2.8185 nM | 0.9023 | 4 |
| Supp. Figure 2d | $ATV^{TfR}$-A 3 months | CSF 1-day post-dose | 2.7283 nM | 0.4010 | 5 |
| Supp. Figure 2d | $ATV^{TfR}$-A 12 months | CSF 1-day post-dose | 2.9310 nM | 0.8122 | 5 |
| Supp. Figure 2d | $ATV^{TfR}$-A 24 months | CSF 1-day post-dose | 2.7112 nM | 0.9162 | 5 |
| Supp. Figure 2d | Control huIgG 1.5 months | CSF 4-days post-dose | 1.6024 nM | 0.9391 | 5 |
| Supp. Figure 2d | Control huIgG 3 months | CSF 4-days post-dose | 1.0346 nM | 0.4547 | 5 |
| Supp. Figure 2d | Control huIgG 12 months | CSF 4-days post-dose | 1.0771 nM | 0.2939 | 5 |
| Supp. Figure 2d | Control huIgG 24 months | CSF 4-days post-dose | 1.5644 nM | 0.7387 | 5 |
| Supp. Figure 2d | $ATV^{TfR}$-A 1.5 months | CSF 4-days post-dose | 1.2657 nM | 0.1919 | 5 |
| Supp. Figure 2d | $ATV^{TfR}$-A 3 months | CSF 4-days post-dose | 1.5452 nM | 0.3837 | 5 |
| Supp. Figure 2d | $ATV^{TfR}$-A 12 months | CSF 4-days post-dose | 1.8205 nM | 0.4217 | 5 |

| | | | | | |
|---|---|---|---|---|---|
| Supp. Figure 2d | $ATV^{TfR}$-A 24 months | CSF 4-days post-dose | 2.4936 nM | 2.2963 | 5 |
| Supp. Figure 2d | Control huIgG 1.5 months | CSF 7-days post-dose | 1.0648 nM | 0.3110 | 5 |
| Supp. Figure 2d | Control huIgG 3 months | CSF 7-days post-dose | 0.9578 nM | 0.2825 | 5 |
| Supp. Figure 2d | Control huIgG 12 months | CSF 7-days post-dose | 0.9863 nM | 0.5693 | 5 |
| Supp. Figure 2d | Control huIgG 24 months | CSF 7-days post-dose | 1.5379 nM | 0.7126 | 5 |
| Supp. Figure 2d | $ATV^{TfR}$-A 1.5 months | CSF 7-days post-dose | 0.6831 nM | 0.1245 | 5 |
| Supp. Figure 2d | $ATV^{TfR}$-A 3 months | CSF 7-days post-dose | 0.8199 nM | 0.2911 | 5 |
| Supp. Figure 2d | $ATV^{TfR}$-A 12 months | CSF 7-days post-dose | 0.9971 nM | 0.4342 | 5 |
| Supp. Figure 2d | $ATV^{TfR}$-A 24 months | CSF 7-days post-dose | 1.0872 nM | 0.3583 | 4 |

**Table S2.** Pairwise statistical comparisons for TfR expression across healthy aging (Fig. 1 and Fig. S1).

| | ANOVAs – Tukey's Multiple Comparisons Test | | | | | | | |
|---|---|---|---|---|---|---|---|---|
| **Figure and Panel** | **Groups** | **Type** | **Unit** | **Group mean diff.** | **df** | **conf. low** | **conf. high** | **Adj P value** |
| Figure 1a | 1.5 months v. 3 months | One-way | (ng/mL)/total protein | 4.3 | 16 | -41 | 49 | 0.9929 |
| Figure 1a | 1.5 months v. 12 months | One-way | (ng/mL)/total protein | 13 | 16 | -32 | 58 | 0.8244 |
| Figure 1a | 1.5 months v. 24 months | One-way | (ng/mL)/total protein | 25 | 16 | -20 | 70 | 0.4229 |
| Figure 1a | 3 months v. 12 months | One-way | (ng/mL)/total protein | 8.8 | 16 | -36 | 54 | 0.9437 |
| Figure 1a | 3 months v. 24 months | One-way | (ng/mL)/total protein | 21 | 16 | -25 | 66 | 0.5763 |
| Figure 1a | 12 months v. 24 months | One-way | (ng/mL)/total protein | 12 | 16 | -33 | 57 | 0.8777 |
| Figure 1c | 1.5 months v. 3 months | One-way | AU/mm$^2$ | 858.2 | 16 | -750.6 | 2467 | 0.4457 |
| Figure 1c | 1.5 months v. 12 months | One-way | AU/mm$^2$ | 2689 | 16 | 1081 | 4298 | 1.05E-03 |
| Figure 1c | 1.5 months v. 24 months | One-way | AU/mm$^2$ | 3355 | 16 | 1746 | 4963 | 1.06E-04 |
| Figure 1c | 3 months v. 12 months | One-way | AU/mm$^2$ | 1831 | 16 | 222.3 | 3440 | 2.31E-02 |
| Figure 1c | 3 months v. 24 months | One-way | AU/mm$^2$ | 2496 | 16 | 887.5 | 4105 | 2.10E-03 |
| Figure 1c | 12 months v. 24 months | One-way | AU/mm$^2$ | 665.2 | 16 | -943.6 | 2274 | 0.6457 |
| Figure 1d | 1.5 months v. 3 months | One-way | (pg/mL)/cell | 0.01287 | 16 | -0.1072 | 0.13.29 | 0.9896 |
| Figure 1d | 1.5 months v. 16 months | One-way | (pg/mL)/cell | 0.1183 | 16 | -0.001803 | 0.2383 | 5.43E-02 |
| Figure 1d | 1.5 months v. 28 months | One-way | (pg/mL)/cell | 0.2526 | 16 | 0.1325 | 0.3726 | 9.57E-05 |
| Figure 1d | 3 months v. 16 months | One-way | (pg/mL)/cell | 0.1054 | 16 | -0.01467 | 0.2254 | 9.60E-02 |
| Figure 1d | 3 months v. 28 months | One-way | (pg/mL)/cell | 0.2397 | 16 | 0.1196 | 0.3597 | 1.71E-04 |
| Figure 1d | 16 months v. 28 months | One-way | (pg/mL)/cell | 0.1343 | 16 | 0.01425 | 0.25.44 | 2.58E-02 |
| Figure 1e | 3 months v. 15 months | One-way | AU/mm$^3$ | 0.01249 | 8 | -10.76 | 12.06 | 0.9855 |
| Figure 1e | 3 months v. 21.5 months | One-way | AU/mm$^3$ | 0.07382 | 8 | -0.9919 | 23.66 | 7.01E-02 |
| Figure 1e | 15 months v. 21.5 months | One-way | AU/mm$^3$ | 0.06133 | 8 | -1.642 | 23.01 | 8.75E-02 |
| Supp. Figure 1a | 0.03 months v. 1.5 months | One-way | (ng/mL)/total protein | 96.05 | 22 | 41.48 | 150.6 | 6.08E-04 |
| Supp. Figure 1a | 0.03 months v. 3-24 months | One-way | (ng/mL)/total protein | 110.1 | 22 | 65.52 | 154.6 | 8.76E-06 |
| Supp. Figure 1a | 1.5 months v. 3-24 months | One-way | (ng/mL)/total protein | 14.03 | 22 | -30.53 | 58.59 | 0.7123 |
| Supp. Figure 1b | 0.03 months v. 1.5 months | One-way | AU/mm$^2$ | 13245 | 20 | 8907 | 17583 | 5.80E-07 |
| Supp. Figure 1b | 0.03 months v. 3-24 months | One-way | AU/mm$^2$ | 15546 | 20 | 11789 | 19303 | 4.27E-09 |
| Supp. Figure 1b | 1.5 months v. 3-24 months | One-way | AU/mm$^2$ | 2301 | 20 | -766.9 | 5368 | 0.1653 |
| Supp. Figure 1c | 1.5 months v. 3 months | One-way | mm$^2$ | 0.03936 | 16 | -0.7715 | 0.8503 | 0.9990 |
| Supp. Figure 1c | 1.5 months v. 12 months | One-way | mm$^2$ | -0.1024 | 16 | -0.9133 | 0.7085 | 0.9832 |

| | | | | | | | | |
|---|---|---|---|---|---|---|---|---|
| Supp. Figure 1c | 1.5 months v. 24 months | One-way | $mm^2$ | -1.111 | 16 | -1.922 | -0.3004 | 6.02E-03 |
| Supp. Figure 1c | 3 months v. 12 months | One-way | $mm^2$ | -0.1418 | 16 | -0.9527 | 0.6691 | 0.9578 |
| Supp. Figure 1c | 3 months v. 24 months | One-way | $mm^2$ | -1.151 | 16 | -1.962 | -0.3397 | 4.54E-03 |
| Supp. Figure 1c | 12 months v. 24 months | One-way | $mm^2$ | -1.009 | 16 | -1.820 | -0.1980 | 1.25E-02 |
| Supp. Figure 1d | 3 months v. 15 months | One-way | $mm^3$ | 0.01249 | 8 | -0.03370 | 0.05868 | 0.7292 |
| Supp. Figure 1d | 3 months v. 21.5 months | One-way | $mm^3$ | 0.07382 | 8 | 0.02393 | 0.1237 | 7.20E-03 |
| Supp. Figure 1d | 15 months v. 21.5 months | One-way | $mm^3$ | 0.06133 | 8 | 0.01145 | 0.1112 | 1.93E-02 |

**Table S3.** Pairwise statistical comparisons for huIgG exposure across healthy aging (Fig. 2-3 and Fig. S2).

| | **ANOVAs – Tukey's Multiple Comparisons Test** | | | | | | | |
|---|---|---|---|---|---|---|---|---|
| **Figure and Panel** | **Groups** | **Type** | **Unit** | **Group mean diff.** | **df** | **conf. low** | **conf. high** | **Adj P value** |
| Figure 2a | Control huIgG (3 months v. 10.5 months) | Two-way | mM | -0.01550 | 38 | -0.2738 | 0.2428 | 0.9882 |
| Figure 2a | Control huIgG (3 months v. 24 months) | Two-way | mM | 0.3538 | 38 | 0.08400 | 0.6236 | 7.67E-03 |
| Figure 2a | Control huIgG (10.5 months v. 24 months) | Two-way | mM | 0.3693 | 38 | 0.1110 | 0.6276 | 3.50E-03 |
| Figure 2a | $ATV^{TfR}$-A (3 months v. 10.5 months) | Two-way | mM | -0.07222 | 38 | -0.3476 | 0.2031 | 0.7992 |
| Figure 2a | $ATV^{TfR}$-A (3 months v. 24 months) | Two-way | mM | 0.1486 | 38 | -0.1375 | 0.4348 | 0.4223 |
| Figure 2a | $ATV^{TfR}$-A (10.5 months v. 24 months) | Two-way | mM | 0.2209 | 38 | -0.0374 | 0.4792 | 0.1064 |
| Figure 2a | $ATV^{TfR}$-B (3 months v. 10.5 months) | Two-way | mM | 0.1265 | 38 | -0.1318 | 0.3848 | 0.4638 |
| Figure 2a | $ATV^{TfR}$-B (3 months v. 24 months) | Two-way | mM | 0.1646 | 38 | -0.1052 | 0.4344 | 0.3077 |
| Figure 2a | $ATV^{TfR}$-B (10.5 months v. 24 months) | Two-way | mM | 0.03814 | 38 | -0.2202 | 0.2964 | 0.9311 |
| Figure 2b | Control huIgG (3 months v. 10.5 months) | Two-way | nM | 0.02641 | 37 | -2.055 | 2.108 | 0.9994 |
| Figure 2b | Control huIgG (3 months v. 24 months) | Two-way | nM | 0.6029 | 37 | -1.703 | 2.909 | 0.8000 |
| Figure 2b | Control huIgG (10.5 months v. 24 months) | Two-way | nM | 0.5764 | 37 | -1.643 | 2.795 | 0.8023 |
| Figure 2b | $ATV^{TfR}$-A (3 months v. 10.5 months) | Two-way | nM | 0.6626 | 37 | -1.556 | 2.882 | 0.7479 |
| Figure 2b | $ATV^{TfR}$-A (3 months v. 24 months) | Two-way | nM | 3.035 | 37 | 0.7294 | 5.341 | 7.40E-03 |
| Figure 2b | $ATV^{TfR}$-A (10.5 months v. 24 months) | Two-way | nM | 2.373 | 37 | 0.2912 | 4.454 | 2.24E-02 |
| Figure 2b | $ATV^{TfR}$-B (3 months v. 10.5 months) | Two-way | nM | 2.420 | 37 | 0.3379 | 4.501 | 1.95E-02 |
| Figure 2b | $ATV^{TfR}$-B (3 months v. 24 months) | Two-way | nM | 3.491 | 37 | 1.317 | 4.501 | 1.00E-03 |
| Figure 2b | $ATV^{TfR}$-B (10.5 months v. 24 months) | Two-way | nM | 1.072 | 37 | -1.010 | 3.153 | 0.4280 |
| Figure 2c | Control huIgG (3 months v. 10.5 months) | Two-way | % brain: plasma | 0.005760 | 38 | -1.341 | 1.352 | 0.9999 |
| Figure 2c | Control huIgG (3 months v. 24 months) | Two-way | % brain: plasma | 0.05446 | 38 | -1.352 | 1.461 | 0.9950 |
| Figure 2c | Control huIgG (10.5 months v. 24 months) | Two-way | % brain: plasma | 0.04870 | 38 | -1.298 | 1.395 | 0.9957 |
| Figure 2c | $ATV^{TfR}$-A (3 months v. 10.5 months) | Two-way | % brain: plasma | 0.2200 | 38 | -1.215 | 1.655 | 0.9260 |
| Figure 2c | $ATV^{TfR}$-A (3 months v. 24 months) | Two-way | % brain: plasma | 0.2364 | 38 | -1.255 | 1.728 | 0.9211 |
| Figure 2c | $ATV^{TfR}$-A (10.5 months v. 24 months) | Two-way | % brain: plasma | 0.01647 | 38 | -1.330 | 1.363 | 0.9995 |
| Figure 2c | $ATV^{TfR}$-B (3 months v. 10.5 months) | Two-way | % brain: plasma | -0.7192 | 38 | -2.066 | 0.6273 | 0.4024 |
| Figure 2c | $ATV^{TfR}$-B (3 months v. 24 months) | Two-way | % brain: plasma | -0.4621 | 38 | -1.869 | 0.9443 | 0.7044 |
| Figure 2c | $ATV^{TfR}$-B (10.5 months v. 24 months) | Two-way | % brain: plasma | 0.2571 | 38 | -1.089 | 1.604 | 0.8877 |
| Figure 2d | Control huIgG (3 months v. 10.5 months) | Two-way | % CSF: plasma | 0.01328 | 32 | -0.1756 | 0.2022 | 0.9836 |
| Figure 2d | Control huIgG (3 months v. 24 months) | Two-way | % CSF: plasma | 0.03045 | 32 | -0.1669 | 0.2278 | 0.9239 |

| | | | | | | | | |
|---|---|---|---|---|---|---|---|---|
| Figure 2d | Control huIgG (10.5 months v. 24 months) | Two-way | % CSF: plasma | 0.01718 | 32 | -0.1717 | 0.2061 | 0.9728 |
| Figure 2d | ATV$^{TfR}$-A (3 months v. 10.5 months) | Two-way | % CSF: plasma | -0.02028 | 32 | -0.2409 | 0.2003 | 0.9722 |
| Figure 2d | ATV$^{TfR}$-A (3 months v. 24 months) | Two-way | % CSF: plasma | 0.01155 | 32 | -0.2432 | 0.2663 | 0.9931 |
| Figure 2d | ATV$^{TfR}$-A (10.5 months v. 24 months) | Two-way | % CSF: plasma | 0.03183 | 32 | -0.1888 | 0.2524 | 0.9331 |
| Figure 2d | ATV$^{TfR}$-B (3 months v. 10.5 months) | Two-way | % CSF: plasma | -0.06562 | 32 | -0.2629 | 0.1317 | 0.6952 |
| Figure 2d | ATV$^{TfR}$-B (3 months v. 24 months) | Two-way | % CSF: plasma | -0.1436 | 32 | -0.3714 | 0.0842 5 | 0.2823 |
| Figure 2d | ATV$^{TfR}$-B (10.5 months v. 24 months) | Two-way | % CSF: plasma | -0.07798 | 32 | -0.3058 | 0.1499 | 0.6806 |
| Figure 3c | Control huIgG (1.5 months v. 3 months) | Two-way | % brain: plasma | 0.003800 | 32 | -0.3700 | 0.3776 | 0.9999 |
| Figure 3c | Control huIgG (1.5 months v. 12 months) | Two-way | % brain: plasma | 0.01180 | 32 | -0.3620 | 0.3856 | 0.9999 |
| Figure 3c | Control huIgG (1.5 months v. 24 months) | Two-way | % brain: plasma | -0.005600 | 32 | -0.3794 | 0.3682 | 0.9999 |
| Figure 3c | Control huIgG (3 months v. 12 months) | Two-way | % brain: plasma | 0.008000 | 32 | -0.3658 | 0.3818 | 0.9999 |
| Figure 3c | Control huIgG (3 months v. 24 months) | Two-way | % brain: plasma | -0.009400 | 32 | -0.3832 | 0.3644 | 0.9999 |
| Figure 3c | Control huIgG (12 months v. 24 months) | Two-way | % brain: plasma | -0.01740 | 32 | -0.3912 | 0.3564 | 0.9999 |
| Figure 3c | ATV$^{TfR}$-A (1.5 months v. 3 months) | Two-way | % brain: plasma | 1.036 | 32 | 0.6620 | 1.410 | 7.98E-09 |
| Figure 3c | ATV$^{TfR}$-A (1.5 months v. 12 months) | Two-way | % brain: plasma | 1.397 | 32 | 1.023 | 1.771 | 4.85E-12 |
| Figure 3c | ATV$^{TfR}$-A (1.5 months v. 24 months) | Two-way | % brain: plasma | 1.319 | 32 | 0.9448 | 1.692 | 2.15E-11 |
| Figure 3c | ATV$^{TfR}$-A (3 months v. 12 months) | Two-way | % brain: plasma | 0.3612 | 32 | - 0.0125 | 0.7350 | 6.42E-02 |
| Figure 3c | ATV$^{TfR}$-A (3 months v. 24 months) | Two-way | % brain: plasma | 0.2828 | 32 | - 0.0909 | 0.6566 | 0.2525 |
| Figure 3c | ATV$^{TfR}$-A (12 months v. 24 months) | Two-way | % brain: plasma | -0.07840 | 32 | -0.4522 | 0.2954 | 0.9970 |
| Figure 3f | Control huIgG (1.5 months v. 3 months) | Two-way | nM | -0.001026 | 28 | - 0.0890 | 0.0869 9 | 0.9999 |
| Figure 3f | Control huIgG (1.5 months v. 12 months) | Two-way | nM | -0.002999 | 28 | - 0.0871 | 0.0811 6 | 0.9999 |
| Figure 3f | Control huIgG (1.5 months v. 24 months) | Two-way | nM | -0.002503 | 28 | - 0.0905 | 0.0855 1 | 0.9999 |
| Figure 3f | Control huIgG (3 months v. 12 months) | Two-way | nM | -0.001974 | 28 | - 0.0792 | 0.0753 3 | 0.9999 |
| Figure 3f | Control huIgG (3 months v. 24 months) | Two-way | nM | -0.001478 | 28 | - 0.0829 | 0.0800 1 | 0.9999 |
| Figure 3f | Control huIgG (12 months v. 24 months) | Two-way | nM | 0.0004960 | 28 | - 0.0768 | 0.0778 0 | >0.9999 |
| Figure 3f | ATV$^{TfR}$-A (1.5 months v. 3 months) | Two-way | nM | 0.04799 | 28 | - 0.0248 | 0.1209 | 0.4079 |
| Figure 3f | ATV$^{TfR}$-A (1.5 months v. 12 months) | Two-way | nM | 0.05620 | 28 | - 0.0166 | 0.1291 | 0.2276 |
| Figure 3f | ATV$^{TfR}$-A (1.5 months v. 24 months) | Two-way | nM | 0.04899 | 28 | - 0.0238 | 0.1219 | 0.3825 |
| Figure 3f | ATV$^{TfR}$-A (3 months v. 12 months) | Two-way | nM | 0.008206 | 28 | - 0.0646 | 0.0810 9 | 0.9999 |
| Figure 3f | ATV$^{TfR}$-A (3 months v. 24 months) | Two-way | nM | 0.001004 | 28 | - 0.0718 | 0.0738 9 | 0.9999 |
| Figure 3f | ATV$^{TfR}$-A (12 months v. 24 months) | Two-way | nM | -0.007202 | 28 | - 0.0800 | 0.0656 8 | 0.9999 |
| Figure 3h | ATV$^{TfR}$-A (1.5 months v. 12 months) | One-way | AU/mm$^2$ | 1217 | 16 | 515.6 | 1919 | 7.33E-04 |
| Figure 3h | ATV$^{TfR}$-A (1.5 months v. 24 months) | One-way | AU/mm$^2$ | 1095 | 16 | 393.7 | 1797 | 1.99E-03 |

| | | | | | | | | |
|---|---|---|---|---|---|---|---|---|
| Figure 3h | ATV$^{TfR}$-A (3 months v. 12 months) | One-way | AU/mm$^2$ | 559.4 | 16 | -142.2 | 1261 | 0.1441 |
| Figure 3h | ATV$^{TfR}$-A (3 months v. 24 months) | One-way | AU/mm$^2$ | 437.6 | 16 | -264.1 | 1139 | 0.3161 |
| Figure 3h | ATV$^{TfR}$-A (12 months v. 24 months) | One-way | AU/mm$^2$ | -121.8 | 16 | -823.5 | 579.8 | 0.9586 |
| Figure 3i | ATV$^{TfR}$-A (1.5 months v. 3 months) | One-way | AU/mm$^2$ | 1010 | 16 | -157.6 | 2177 | 0.1025 |
| Figure 3i | ATV$^{TfR}$-A (1.5 months v. 12 months) | One-way | AU/mm$^2$ | 1735 | 16 | 567.5 | 2902 | 3.07E-03 |
| Figure 3i | ATV$^{TfR}$-A (1.5 months v. 24 months) | One-way | AU/mm$^2$ | 1508 | 16 | 340.5 | 2675 | 9.52E-03 |
| Figure 3i | ATV$^{TfR}$-A (3 months v. 12 months) | One-way | AU/mm$^2$ | 725.1 | 16 | -442.2 | 1892 | 0.3193 |
| Figure 3i | ATV$^{TfR}$-A (3 months v. 24 months) | One-way | AU/mm$^2$ | 498.1 | 16 | -669.2 | 1665 | 0.6232 |
| Figure 3i | ATV$^{TfR}$-A (12 months v. 24 months) | One-way | AU/mm$^2$ | -227.0 | 16 | -1394 | 940.3 | 0.9434 |
| Figure 3j | ATV$^{TfR}$-A (1.5 months v. 3 months) | One-way | AU/mm$^2$ | 610.2 | 16 | -69.41 | 1290 | 8.65E-02 |
| Figure 3j | ATV$^{TfR}$-A (1.5 months v. 12 months) | One-way | AU/mm$^2$ | 1163 | 16 | 483.4 | 1843 | 8.38E-04 |
| Figure 3j | ATV$^{TfR}$-A (1.5 months v. 24 months) | One-way | AU/mm$^2$ | 1060 | 16 | 380.1 | 1739 | 2.01E-03 |
| Figure 3j | ATV$^{TfR}$-A (3 months v. 12 months) | One-way | AU/mm$^2$ | 552.8 | 16 | -126.8 | 1232 | 0.1331 |
| Figure 3j | ATV$^{TfR}$-A (3 months v. 24 months) | One-way | AU/mm$^2$ | 449.6 | 16 | -230.0 | 1129 | 0.2698 |
| Figure 3j | ATV$^{TfR}$-A (12 months v. 24 months) | One-way | AU/mm$^2$ | -103.3 | 16 | -782.9 | 576.3 | 0.9715 |
| Supp. Figure 2a | Control huIgG (3 months v. 10.5 months) | Two-way | nM | 0.1162 | 32 | -0.8039 | 1.036 | 0.9483 |
| Supp. Figure 2a | Control huIgG (3 months v. 24 months) | Two-way | nM | 0.9385 | 32 | -0.0225 | 1.900 | 5.67E-02 |
| Supp. Figure 2a | Control huIgG (10.5 months v. 24 months) | Two-way | nM | 0.8223 | 32 | -0.0977 | 1.742 | 8.72E-02 |
| Supp. Figure 2a | ATV$^{TfR}$-A (3 months v. 10.5 months) | Two-way | nM | -0.2308 | 32 | -1.305 | 0.8436 | 0.8583 |
| Supp. Figure 2a | ATV$^{TfR}$-A (3 months v. 24 months) | Two-way | nM | 0.8024 | 32 | -0.4383 | 2.043 | 0.2648 |
| Supp. Figure 2a | ATV$^{TfR}$-A (10.5 months v. 24 months) | Two-way | nM | 1.033 | 32 | -0.0412 | 2.108 | 6.14E-02 |
| Supp. Figure 2a | ATV$^{TfR}$-B (3 months v. 10.5 months) | Two-way | nM | -0.05905 | 32 | -1.020 | 0.9020 | 0.9875 |
| Supp. Figure 2a | ATV$^{TfR}$-B (3 months v. 24 months) | Two-way | nM | -0.009280 | 32 | -1.119 | 1.100 | 0.9997 |
| Supp. Figure 2a | ATV$^{TfR}$-B (10.5 months v. 24 months) | Two-way | nM | 0.04977 | 32 | -1.060 | 1.159 | 0.9933 |
| Supp. Figure 2b | 3 months v. 10.5 months | One-way | % brain: plasma | 0.005760 | 12 | -0.0516 | 0.06312 | 0.9613 |
| Supp. Figure 2b | 3 months v. 24 months | One-way | % brain: plasma | 0.03230 | 12 | -0.0312 | 0.09584 | 0.3933 |
| Supp. Figure 2b | 10.5 months v. 24 months | One-way | % brain: plasma | 0.02654 | 12 | -0.0346 | 0.08769 | 0.4991 |
| Supp. Figure 2c | Control huIgG (0.03 months v. 1.5 months) | Two-way | nM | 3.523 | 45 | -0.3647 | 7.410 | 9.58E-02 |
| Supp. Figure 2c | Control huIgG (0.03 months v. 3-24 months) | Two-way | nM | 3.391 | 45 | 0.2167 | 6.565 | 3.01E-02 |
| Supp. Figure 2c | Control huIgG (1.5 months v. 3-24 months) | Two-way | nM | -0.1319 | 45 | -3.306 | 3.042 | 0.9999 |
| Supp. Figure 2c | ATV$^{TfR}$-A (0.03 months v. 1.5 months) | Two-way | nM | 15.13 | 45 | 11.40 | 18.85 | 2.69E-13 |
| Supp. Figure 2c | ATV$^{TfR}$-A (0.03 months v. 3-24 months) | Two-way | nM | 19.14 | 45 | 16.17 | 22.11 | 2.40E-13 |
| Supp. Figure 2c | ATV$^{TfR}$-A (1.5 months v. 3-24 months) | Two-way | nM | 4.017 | 45 | 0.8431 | 7.191 | 5.99E-03 |

| | | | | | | | | |
|---|---|---|---|---|---|---|---|---|
| Supp. Figure 2e | Control huIgG (1.5 months v. 3 months) | Two-way | nM | 0.1700 | 30 | -2.485 | 2.825 | 0.9999 |
| Supp. Figure 2e | Control huIgG (1.5 months v. 12 months) | Two-way | nM | -0.2583 | 30 | -2.913 | 2.396 | 0.9999 |
| Supp. Figure 2e | Control huIgG (1.5 months v. 24 months) | Two-way | nM | -0.8325 | 30 | -3.648 | 1.983 | 0.9764 |
| Supp. Figure 2e | Control huIgG (3 months v. 12 months) | Two-way | nM | -0.4283 | 30 | -3.083 | 2.226 | 0.9994 |
| Supp. Figure 2e | Control huIgG (3 months v. 24 months) | Two-way | nM | -1.002 | 30 | -3.818 | 1.813 | 0.9374 |
| Supp. Figure 2e | Control huIgG (12 months v. 24 months) | Two-way | nM | -0.5741 | 30 | -3.390 | 2.242 | 0.9973 |
| Supp. Figure 2e | $ATV^{TfR}$-A (1.5 months v. 3 months) | Two-way | nM | 0.09027 | 30 | -2.725 | 2.906 | 0.9999 |
| Supp. Figure 2e | $ATV^{TfR}$-A (1.5 months v. 12 months) | Two-way | nM | -0.1124 | 30 | -2.928 | 2.703 | 0.9999 |
| Supp. Figure 2e | $ATV^{TfR}$-A (1.5 months v. 24 months) | Two-way | nM | 0.1073 | 30 | -2.708 | 2.923 | 0.9999 |
| Supp. Figure 2e | $ATV^{TfR}$-A (3 months v. 12 months) | Two-way | nM | -0.2027 | 30 | -2.857 | 2.452 | 0.9999 |
| Supp. Figure 2e | $ATV^{TfR}$-A (3 months v. 24 months) | Two-way | nM | 0.01704 | 30 | -2.638 | 2.672 | >0.9999 |
| Supp. Figure 2e | $ATV^{TfR}$-A (12 months v. 24 months) | Two-way | nM | 0.2197 | 30 | -2.435 | 2.874 | >0.9999 |
| Supp. Figure 2f | Control huIgG (1.5 months v. 3 months) | Two-way | % CSF: plasma | 0.05700 | 30 | -0.3192 | 0.4332 | 0.9996 |
| Supp. Figure 2f | Control huIgG (1.5 months v. 12 months) | Two-way | % CSF: plasma | 0.01480 | 30 | -0.3614 | 0.3910 | 0.9999 |
| Supp. Figure 2f | Control huIgG (1.5 months v. 24 months) | Two-way | % CSF: plasma | -0.06105 | 30 | -0.4601 | 0.3380 | 0.9995 |
| Supp. Figure 2f | Control huIgG (3 months v. 12 months) | Two-way | % CSF: plasma | -0.04220 | 30 | -0.4184 | 0.3340 | 0.9999 |
| Supp. Figure 2f | Control huIgG (3 months v. 24 months) | Two-way | % CSF: plasma | -0.1181 | 30 | -0.5171 | 0.2810 | 0.9763 |
| Supp. Figure 2f | Control huIgG (12 months v. 24 months) | Two-way | % CSF: plasma | -0.07585 | 30 | -0.4749 | 0.3232 | 0.9983 |
| Supp. Figure 2f | $ATV^{TfR}$-A (1.5 months v. 3 months) | Two-way | % CSF: plasma | 0.1552 | 30 | -0.2439 | 0.5543 | 0.9043 |
| Supp. Figure 2f | $ATV^{TfR}$-A (1.5 months v. 12 months) | Two-way | % CSF: plasma | 0.1348 | 30 | -0.2643 | 0.5339 | 0.9520 |
| Supp. Figure 2f | $ATV^{TfR}$-A (1.5 months v. 24 months) | Two-way | % CSF: plasma | 0.1530 | 30 | -0.2461 | 0.5521 | 0.9105 |
| Supp. Figure 2f | $ATV^{TfR}$-A (3 months v. 12 months) | Two-way | % CSF: plasma | -0.02040 | 30 | -0.3966 | 0.3558 | 0.9999 |
| Supp. Figure 2f | $ATV^{TfR}$-A (3 months v. 24 months) | Two-way | % CSF: plasma | -0.002200 | 30 | -0.3784 | 0.3740 | >0.9999 |
| Supp. Figure 2f | $ATV^{TfR}$-A (12 months v. 24 months) | Two-way | % CSF: plasma | 0.01820 | 30 | -0.3580 | 0.3944 | 0.9999 |
| Supp. Figure 2g | $ATV^{TfR}$-A (0.03 months v. 1.5 months) | One-way | AU/mm$^2$ | 12403 | 20 | 10272 | 14534 | 1.02E-11 |
| Supp. Figure 2g | $ATV^{TfR}$-A (0.03 months v. 3-24 months) | One-way | AU/mm$^2$ | 13393 | 20 | 11547 | 15239 | 1.39E-13 |
| Supp. Figure 2g | $ATV^{TfR}$-A (1.5 months v. 3-24 months) | One-way | AU/mm$^2$ | 990.1 | 20 | -517.0 | 2497 | 0.2440 |

**Table S4.** Pairwise statistical comparisons for TfR expression and huIgG exposure in disease (Fig. 4 and Fig. S3). Values below the lower limit of quantification are indicated as 'BLLOQ'.

| | **ANOVAs – Tukey's Multiple Comparisons Test** | | | | | | | |
|---|---|---|---|---|---|---|---|---|
| **Figure and Panel** | **Groups** | **Type** | **Unit** | **Group mean diff.** | **df** | **conf. low** | **conf. high** | **Adj P value** |
| Figure 4a | 1.5 months v. 3 months | One-way | %Ab plaque: tissue | -0.02193 | 6 | -2.499 | 2.455 | 0.9995 |
| Figure 4a | 1.5 months v. 10.5 months | One-way | %Ab plaque: tissue | -3.766 | 6 | -6.243 | -1.289 | 8.23E-03 |
| Figure 4a | 3 months v. 10.5 months | One-way | %Ab plaque: tissue | -3.744 | 6 | -6.221 | -1.267 | 8.46E-03 |
| Figure 4b | TfR$^{mu/hu}$-KI (1.5 months v. 3 months) | Two-way | (ng/mL)/ total protein | -15.64 | 24 | -73.58 | 42.30 | 0.9578 |
| Figure 4b | TfR$^{mu/hu}$-KI (1.5 months v. 10.5 months) | Two-way | (ng/mL)/ total protein | -2.839 | 24 | -60.78 | 55.10 | 0.9999 |
| Figure 4b | TfR$^{mu/hu}$-KI (3 months v. 10.5 months) | Two-way | (ng/mL)/ total protein | 12.80 | 24 | -45.14 | 70.74 | 0.9821 |
| Figure 4b | 5xFAD; TfR$^{mu/hu}$-KI (1.5 months v. 3 months) | Two-way | (ng/mL)/ total protein | 7.525 | 24 | -50.41 | 65.46 | 0.9984 |
| Figure 4b | 5xFAD; TfR$^{mu/hu}$-KI (1.5 months v. 10.5 months) | Two-way | (ng/mL)/ total protein | 33.38 | 24 | -24.56 | 91.32 | 0.4953 |
| Figure 4b | 5xFAD; TfR$^{mu/hu}$-KI (3 months v. 10.5 months) | Two-way | (ng/mL)/ total protein | 25.86 | 24 | -32.08 | 83.80 | 0.7381 |
| Figure 4b | 1.5 months (TfR$^{mu/hu}$-KI v. 5xFAD; TfR$^{mu/hu}$-KI) | Two-way | (ng/mL)/ total protein | -17.64 | 24 | -75.58 | 40.30 | 0.9314 |
| Figure 4b | 3 months (TfR$^{mu/hu}$-KI v. 5xFAD; TfR$^{mu/hu}$-KI) | Two-way | (ng/mL)/ total protein | 5.526 | 24 | -52.41 | 63.46 | 0.9996 |
| Figure 4b | 10.5 months (TfR$^{mu/hu}$-KI v. 5xFAD; TfR$^{mu/hu}$-KI) | Two-way | (ng/mL)/ total protein | 18.58 | 24 | -39.36 | 76.52 | 0.9161 |
| Figure 4c | TfR$^{mu/hu}$-KI (1.5 months v. 3 months) | Two-way | (pg/mL)/cell | -0.06277 | 18 | -0.340 | 0.2150 | 0.9771 |
| Figure 4c | TfR$^{mu/hu}$-KI (1.5 months v. 10.5 months) | Two-way | (pg/mL)/cell | 0.1330 | 18 | -0.144 | 0.4107 | 0.6560 |
| Figure 4c | TfR$^{mu/hu}$-KI (3 months v. 10.5 months) | Two-way | (pg/mL)/cell | 0.1957 | 18 | -0.082 | 0.4735 | 0.2683 |
| Figure 4c | 5xFAD; TfR$^{mu/hu}$-KI (1.5 months v. 3 months) | Two-way | (pg/mL)/cell | 0.3738 | 18 | 0.09600 | 0.6516 | 5.14E-03 |
| Figure 4c | 5xFAD; TfR$^{mu/hu}$-KI (1.5 months v. 10.5 months) | Two-way | (pg/mL)/cell | 0.1099 | 18 | -0.167 | 0.3877 | 0.8034 |
| Figure 4c | 5xFAD; TfR$^{mu/hu}$-KI (3 months v. 10.5 months) | Two-way | (pg/mL)/cell | -0.2639 | 18 | -0.541 | 0.01388 | 6.82E-02 |
| Figure 4c | 1.5 months (TfR$^{mu/hu}$-KI v. 5xFAD; TfR$^{mu/hu}$-KI) | Two-way | (pg/mL)/cell | -0.2724 | 18 | -0.550 | 0.005380 | 5.64E-02 |
| Figure 4c | 3 months (TfR$^{mu/hu}$-KI v. 5xFAD; TfR$^{mu/hu}$-KI) | Two-way | (pg/mL)/cell | 0.1641 | 18 | -0.113 | 0.4419 | 0.4454 |
| Figure 4c | 10.5 months (TfR$^{mu/hu}$-KI v. 5xFAD; TfR$^{mu/hu}$-KI) | Two-way | (pg/mL)/cell | -0.2955 | 18 | -0.573 | -0.017 | 3.33E-02 |
| Figure 4d | Control huIgG, 5xFAD (1.5 months v. 3 months) | Two-way | nM | -0.3402 | 30 | -0.900 | 0.2196 | 0.4730 |
| Figure 4d | Control huIgG, 5xFAD (1.5 months v. 10.5 months) | Two-way | nM | -0.4270 | 30 | -0.986 | 0.1328 | 0.2511 |
| Figure 4d | Control huIgG, 5xFAD (3 months v. 10.5 months) | Two-way | nM | -0.08680 | 30 | -0.646 | 0.4730 | 0.9998 |
| Figure 4d | ATV$^{TfR}$-A, TfR$^{mu/hu}$-KI (1.5 months v. 3 months) | Two-way | nM | -0.09272 | 30 | -0.526 | 0.3409 | 0.9981 |
| Figure 4d | ATV$^{TfR}$-A, TfR$^{mu/hu}$-KI (1.5 months v. 10.5 months) | Two-way | nM | -0.1241 -0.03141 | 30 | -0.557 | 0.3095 | 0.9872 |
| Figure 4d | ATV$^{TfR}$-A, TfR$^{mu/hu}$-KI (3 months v. 10.5 months) | Two-way | nM | -0.03141 | 30 | -0.465 | 0.4022 | 0.9999 |
| Figure 4d | ATV$^{TfR}$-A, 5xFAD (1.5 months v. 3 months) | Two-way | nM | 0.04867 | 30 | -0.384 | 0.4823 | 0.9999 |

| | | | | | | | | |
|---|---|---|---|---|---|---|---|---|
| Figure 4d | ATV$^{TfR}$-A, 5xFAD (1.5 months v. 10.5 months) | Two-way | nM | 0.03854 | 30 | -0.395 | 0.4721 | 0.9999 |
| Figure 4d | ATV$^{TfR}$-A, 5xFAD (3 months v. 10.5 months) | Two-way | nM | -0.01013 | 30 | -0.443 | 0.4235 | 0.9999 |
| Figure 4d | ATV$^{TfR}$-A, 1.5 months (TfR$^{mu/hu}$-KI v. 5xFAD) | Two-way | nM | 0.005505 | 30 | -0.428 | 0.4391 | >0.9999 |
| Figure 4d | ATV$^{TfR}$-A, 3 months (TfR$^{mu/hu}$-KI v. 5xFAD) | Two-way | nM | -0.1359 | 30 | -0.569 | 0.2977 | 0.9775 |
| Figure 4d | ATV$^{TfR}$-A, 10.5 months (TfR$^{mu/hu}$-KI v. 5xFAD) | Two-way | nM | -0.1572 | 30 | -0.590 | 0.2764 | 0.9480 |
| Figure 4e | Control huIgG, 5xFAD (1.5 months v. 3 months) | Two-way | nM | -0.1337 | 30 | -4.394 | 4.126 | 0.9999 |
| Figure 4e | Control huIgG, 5xFAD (1.5 months v. 10.5 months) | Two-way | nM | -0.1122 | 30 | -4.372 | 4.148 | 0.9999 |
| Figure 4e | Control huIgG, 5xFAD (3 months v. 10.5 months) | Two-way | nM | 0.02148 | 30 | -4.239 | 4.281 | >0.9999 |
| Figure 4e | ATV$^{TfR}$-A, TfR$^{mu/hu}$-KI (1.5 months v. 3 months) | Two-way | nM | 2.739 | 30 | -0.560 | 6.039 | 0.1665 |
| Figure 4e | ATV$^{TfR}$-A, TfR$^{mu/hu}$-KI (1.5 months v. 10.5 months) | Two-way | nM | 4.492 | 30 | 1.192 | 7.791 | 2.39E-03 |
| Figure 4e | ATV$^{TfR}$-A, TfR$^{mu/hu}$-KI (3 months v. 10.5 months) | Two-way | nM | 1.753 | 30 | -1.547 | 5.052 | 0.6984 |
| Figure 4e | ATV$^{TfR}$-A, 5xFAD (1.5 months v. 3 months) | Two-way | nM | 2.418 | 30 | -0.881 | 5.718 | 0.2974 |
| Figure 4e | ATV$^{TfR}$-A, 5xFAD (1.5 months v. 10.5 months) | Two-way | nM | 3.424 | 30 | 0.1242 | 6.724 | 3.73E-02 |
| Figure 4e | ATV$^{TfR}$-A, 5xFAD (3 months v. 10.5 months) | Two-way | nM | 1.006 | 30 | -2.294 | 4.306 | 0.9810 |
| Figure 4e | ATV$^{TfR}$-A, 1.5 months (TfR$^{mu/hu}$-KI v. 5xFAD) | Two-way | nM | -1.037 | 30 | -4.336 | 2.263 | 0.9772 |
| Figure 4e | ATV$^{TfR}$-A, 3 months (TfR$^{mu/hu}$-KI v. 5xFAD) | Two-way | nM | -0.7154 | 30 | -4.015 | 2.584 | 0.9980 |
| Figure 4e | ATV$^{TfR}$-A, 10.5 months (TfR$^{mu/hu}$-KI v. 5xFAD) | Two-way | nM | 0.03120 | 30 | -3.269 | 3.331 | >0.9999 |
| Figure 4f | Control huIgG, 5xFAD (1.5 months v. 3 months) | - | BLLOQ | - | - | - | - | - |
| Figure 4f | Control huIgG, 5xFAD (1.5 months v. 10.5 months) | - | BLLOQ | - | - | - | - | - |
| Figure 4f | Control huIgG, 5xFAD (3 months v. 10.5 months) | - | BLLOQ | - | - | - | - | - |
| Figure 4f | ATV$^{TfR}$-A, TfR$^{mu/hu}$-KI (1.5 months v. 3 months) | Two-way | nM | 0.008480 | 24 | -0.163 | 0.1806 | 0.9999 |
| Figure 4f | ATV$^{TfR}$-A, TfR$^{mu/hu}$-KI (1.5 months v. 10.5 months) | Two-way | nM | 0.03505 0.02657 | 24 | -0.137 | 0.2072 | 0.9876 |
| Figure 4f | ATV$^{TfR}$-A, TfR$^{mu/hu}$-KI (3 months v. 10.5 months) | Two-way | nM | 0.02657 | 24 | -0.145 | 0.1987 | 0.9965 |
| Figure 4f | ATV$^{TfR}$-A, 5xFAD (1.5 months v. 3 months) | Two-way | nM | 0.08509 | 24 | -0.087 | 0.2572 | 0.6503 |
| Figure 4f | ATV$^{TfR}$-A, 5xFAD (1.5 months v. 10.5 months) | Two-way | nM | 0.1258 | 24 | -0.046 | 0.2979 | 0.2491 |
| Figure 4f | ATV$^{TfR}$-A, 5xFAD (3 months v. 10.5 months) | Two-way | nM | 0.04068 | 24 | -0.131 | 0.2128 | 0.9760 |
| Figure 4f | ATV$^{TfR}$-A, 1.5 months (TfR$^{mu/hu}$-KI v. 5xFAD) | Two-way | nM | 0.04936 | 24 | -0.122 | 0.2215 | 0.9461 |
| Figure 4f | ATV$^{TfR}$-A, 3 months (TfR$^{mu/hu}$-KI v. 5xFAD) | Two-way | nM | -0.02725 | 24 | -0.199 | 0.1449 | 0.9961 |
| Figure 4f | ATV$^{TfR}$-A, 10.5 months (TfR$^{mu/hu}$-KI v. 5xFAD) | Two-way | nM | -0.04136 | 24 | -0.213 | 0.1308 | 0.9742 |
| Figure 4g | Control huIgG, 5xFAD (1.5 months v. 3 months) | - | BLLOQ | - | - | - | - | - |
| Figure 4g | Control huIgG, 5xFAD (1.5 months v. 10.5 months) | - | BLLOQ | - | - | - | - | - |
| Figure 4g | Control huIgG, 5xFAD (3 months v. 10.5 months) | - | BLLOQ | - | - | - | - | - |
| Figure 4g | ATV$^{TfR}$-A, TfR$^{mu/hu}$-KI (1.5 months v. 3 months) | Two-way | nM | 0.05064 | 24 | -0.008 | 0.1094 | 0.1205 |

| | | | | | | | | |
|---|---|---|---|---|---|---|---|---|
| Figure 4g | ATV$^{TfR}$-A, TfR$^{mu/hu}$-KI (1.5 months v. 10.5 months) | Two-way | nM | 0.1084 | 24 | 0.04956 | 0.1672 | 9.54E-05 |
| Figure 4g | ATV$^{TfR}$-A, TfR$^{mu/hu}$-KI (3 months v. 10.5 months) | Two-way | nM | 0.05772 | 24 | -0.001 | 0.1165 | 5.65E-02 |
| Figure 4g | ATV$^{TfR}$-A, 5xFAD (1.5 months v. 3 months) | Two-way | nM | 0.05773 | 24 | -0.001 | 0.1165 | 5.64E-02 |
| Figure 4g | ATV$^{TfR}$-A, 5xFAD (1.5 months v. 10.5 months) | Two-way | nM | 0.08782 | 24 | 0.02902 | 0.1466 | 1.37E-03 |
| Figure 4g | ATV$^{TfR}$-A, 5xFAD (3 months v. 10.5 months) | Two-way | nM | 0.03009 | 24 | -0.026 | 0.09096 | 0.5505 |
| Figure 4g | ATV$^{TfR}$-A, 1.5 months (TfR$^{mu/hu}$-KI v. 5xFAD) | Two-way | nM | 0.005028 | 24 | -0.053 | 0.06383 | 0.9997 |
| Figure 4g | ATV$^{TfR}$-A, 3 months (TfR$^{mu/hu}$-KI v. 5xFAD) | Two-way | nM | -0.002060 | 24 | -0.060 | 0.05675 | 0.9999 |
| Figure 4g | ATV$^{TfR}$-A, 10.5 months (TfR$^{mu/hu}$-KI v. 5xFAD) | Two-way | nM | 0.02557 | 24 | -0.033 | 0.08437 | 0.7580 |
| Figure 4h | Control huIgG, 5xFAD (1.5 months v. 3 months) | Two-way | nM | 0.7254 | 30 | -4.662 | 6.112 | 0.9999 |
| Figure 4h | Control huIgG, 5xFAD (1.5 months v. 10.5 months) | Two-way | nM | 1.480 | 30 | -3.907 | 6.867 | 0.9901 |
| Figure 4h | Control huIgG, 5xFAD (3 months v. 10.5 months) | Two-way | nM | 0.7549 | 30 | -4.632 | 6.142 | 0.9999 |
| Figure 4h | ATV$^{TfR}$-A, TfR$^{mu/hu}$-KI (1.5 months v. 3 months) | Two-way | nM | 2.797 | 30 | -1.376 | 6.969 | 0.4092 |
| Figure 4h | ATV$^{TfR}$-A, TfR$^{mu/hu}$-KI (1.5 months v. 10.5 months) | Two-way | nM | 2.622 | 30 | -1.550 | 6.795 | 0.4934 |
| Figure 4h | ATV$^{TfR}$-A, TfR$^{mu/hu}$-KI (3 months v. 10.5 months) | Two-way | nM | -0.1743 | 30 | -4.347 | 3.998 | 0.9999 |
| Figure 4h | ATV$^{TfR}$-A, 5xFAD (1.5 months v. 3 months) | Two-way | nM | -0.1255 | 30 | -4.298 | 4.047 | 0.9999 |
| Figure 4h | ATV$^{TfR}$-A, 5xFAD (1.5 months v. 10.5 months) | Two-way | nM | 0.1304 | 30 | -4.042 | 4.303 | 0.9999 |
| Figure 4h | ATV$^{TfR}$-A, 5xFAD (3 months v. 10.5 months) | Two-way | nM | 0.2559 | 30 | -3.917 | 4.429 | 0.9999 |
| Figure 4h | ATV$^{TfR}$-A, 1.5 months (TfR$^{mu/hu}$-KI v. 5xFAD) | Two-way | nM | -2.912 | 30 | -7.085 | 1.260 | 0.3571 |
| Figure 4h | ATV$^{TfR}$-A, 3 months (TfR$^{mu/hu}$-KI v. 5xFAD) | Two-way | nM | 0.009641 | 30 | -4.163 | 4.182 | >0.9999 |
| Figure 4h | ATV$^{TfR}$-A, 10.5 months (TfR$^{mu/hu}$-KI v. 5xFAD) | Two-way | nM | -0.4205 | 30 | -4.593 | 3.752 | 0.9999 |
| Figure 4i | Control huIgG, 5xFAD (1.5 months v. 3 months) | Two-way | % brain: plasma | 0.02693 | 30 | -0.361 | 0.4155 | 0.9999 |
| Figure 4i | Control huIgG, 5xFAD (1.5 months v. 10.5 months) | Two-way | % brain: plasma | 0.03470<br>0.007772 | 30 | -0.353 | 0.4233 | 0.9999 |
| Figure 4i | Control huIgG, 5xFAD (3 months v. 10.5 months) | Two-way | % brain: plasma | 0.007772 | 30 | -0.380 | 0.3963 | 0.9999 |
| Figure 4i | ATV$^{TfR}$-A, TfR$^{mu/hu}$-KI (1.5 months v. 3 months) | Two-way | % brain: plasma | 0.4932 | 30 | 0.1922 | 0.7942 | 1.91E-04 |
| Figure 4i | ATV$^{TfR}$-A, TfR$^{mu/hu}$-KI (1.5 months v. 10.5 months) | Two-way | % brain: plasma | 0.7107 | 30 | 0.4098 | 1.012 | 2.86E-07 |
| Figure 4i | ATV$^{TfR}$-A, TfR$^{mu/hu}$-KI (3 months v. 10.5 months) | Two-way | % brain: plasma | 0.2175 | 30 | -0.083 | 0.5185 | 0.3138 |
| Figure 4i | ATV$^{TfR}$-A, 5xFAD (1.5 months v. 3 months) | Two-way | % brain: plasma | 0.1946 | 30 | -0.106 | 0.4956 | 0.4562 |
| Figure 4i | ATV$^{TfR}$-A, 5xFAD (1.5 months v. 10.5 months) | Two-way | % brain: plasma | 0.3414 | 30 | 0.04044 | 0.6424 | 1.71E-02 |
| Figure 4i | ATV$^{TfR}$-A, 5xFAD (3 months v. 10.5 months) | Two-way | % brain: plasma | 0.1468 | 30 | -0.154 | 0.4478 | 0.7822 |
| Figure 4i | ATV$^{TfR}$-A, 1.5 months (TfR$^{mu/hu}$-KI v. 5xFAD) | Two-way | % brain: plasma | -0.1613 | 30 | -0.462 | 0.1397 | 0.6890 |
| Figure 4i | ATV$^{TfR}$-A, 3 months (TfR$^{mu/hu}$-KI v. 5xFAD) | Two-way | % brain: plasma | 0.1374 | 30 | -0.163 | 0.4383 | 0.8360 |
| Figure 4i | ATV$^{TfR}$-A, 10.5 months (TfR$^{mu/hu}$-KI v. 5xFAD) | Two-way | % brain: plasma | 0.2080 | 30 | -0.092 | 0.5090 | 0.3694 |
| Figure 4j | Control huIgG, 5xFAD (1.5 months v. 3 months) | Two-way | % CSF: plasma | 0.1054 | 30 | -0.594 | 0.8054 | 0.9998 |

| | | | | | | | | |
|---|---|---|---|---|---|---|---|---|
| Figure 4j | Control huIgG, 5xFAD (1.5 months v. 10.5 months) | Two-way | % CSF: plasma | 0.1671 | 30 | - 0.532 | 0.867 1 | 0.9961 |
| Figure 4j | Control huIgG, 5xFAD (3 months v. 10.5 months) | Two-way | % CSF: plasma | 0.06175 | 30 | - 0.638 | 0.761 8 | 0.9999 |
| Figure 4j | ATV$^{TfR}$-A, TfR$^{mu/hu}$-KI (1.5 months v. 3 months) | Two-way | % CSF: plasma | 0.4141 | 30 | - 0.128 | 0.956 3 | 0.2498 |
| Figure 4j | ATV$^{TfR}$-A, TfR$^{mu/hu}$-KI (1.5 months v. 10.5 months) | Two-way | % CSF: plasma | 0.4076 | 30 | - 0.134 | 0.949 8 | 0.2676 |
| Figure 4j | ATV$^{TfR}$-A, TfR$^{mu/hu}$-KI (3 months v. 10.5 months) | Two-way | % CSF: plasma | -0.006518 | 30 | - 0.548 | 0.535 7 | >0.9999 |
| Figure 4j | ATV$^{TfR}$-A, 5xFAD (1.5 months v. 3 months) | Two-way | % CSF: plasma | -0.05849 | 30 | - 0.600 | 0.483 7 | 0.9999 |
| Figure 4j | ATV$^{TfR}$-A, 5xFAD (1.5 months v. 10.5 months) | Two-way | % CSF: plasma | -0.02781 | 30 | - 0.570 | 0.514 4 | 0.9999 |
| Figure 4j | ATV$^{TfR}$-A, 5xFAD (3 months v. 10.5 months) | Two-way | % CSF: plasma | 0.03068 | 30 | - 0.511 | 0.572 9 | 0.9999 |
| Figure 4j | ATV$^{TfR}$-A, 1.5 months (TfR$^{mu/hu}$-KI v. 5xFAD) | Two-way | % CSF: plasma | -0.3923 | 30 | - 0.934 | 0.149 9 | 0.3124 |
| Figure 4j | ATV$^{TfR}$-A, 3 months (TfR$^{mu/hu}$-KI v. 5xFAD) | Two-way | % CSF: plasma | 0.08025 | 30 | - 0.462 | 0.622 5 | 0.9998 |
| Figure 4j | ATV$^{TfR}$-A, 10.5 months (TfR$^{mu/hu}$-KI v. 5xFAD) | Two-way | % CSF: plasma | 0.04306 | 30 | - 0.499 | 0.585 3 | 0.9999 |
| Figure 4k | TfR$^{mu/hu}$-KI (1.5 months v. 3 months) | Two-way | % brain: plasma | 0.03003 | 24 | - 0.330 | 0.390 8 | 0.9998 |
| Figure 4k | TfR$^{mu/hu}$-KI (1.5 months v. 10.5 months) | Two-way | % brain: plasma | 0.06617 | 24 | - 0.294 | 0.427 0 | 0.9922 |
| Figure 4k | TfR$^{mu/hu}$-KI (3 months v. 10.5 months) | Two-way | % brain: plasma | 0.03614 | 24 | - 0.324 | 0.397 0 | 0.9995 |
| Figure 4k | 5xFAD; TfR$^{mu/hu}$-KI (1.5 months v. 3 months) | Two-way | % brain: plasma | 0.2255 | 24 | - 0.135 | 0.586 3 | 0.4080 |
| Figure 4k | 5xFAD; TfR$^{mu/hu}$-KI (1.5 months v. 10.5 months) | Two-way | % brain: plasma | 0.2308 | 24 | - 0.130 | 0.591 6 | 0.3832 |
| Figure 4k | 5xFAD; TfR$^{mu/hu}$-KI (3 months v. 10.5 months) | Two-way | % brain: plasma | 0.005297 | 24 | - 0.355 | 0.366 1 | 0.9999 |
| Figure 4k | 1.5 months (TfR$^{mu/hu}$-KI v. 5xFAD; TfR$^{mu/hu}$-KI) | Two-way | % brain: plasma | -0.1654 | 24 | - 0.526 | 0.195 4 | 0.7165 |
| Figure 4k | 3 months (TfR$^{mu/hu}$-KI v. 5xFAD; TfR$^{mu/hu}$-KI) | Two-way | % brain: plasma | 0.03003 | 24 | - 0.330 | 0.390 8 | 0.9998 |
| Figure 4k | 10.5 months (TfR$^{mu/hu}$-KI v. 5xFAD; TfR$^{mu/hu}$-KI) | Two-way | % brain: plasma | -0.0008094 | 24 | - 0.361 | 0.360 0 | 0.9999 |
| Figure 4l | TfR$^{mu/hu}$-KI (1.5 months v. 3 months) | Two-way | % CSF: plasma | -0.01898 | 24 | - 0.168 | 0.130 7 | 0.9986 |
| Figure 4l | TfR$^{mu/hu}$-KI (1.5 months v. 10.5 months) | Two-way | % CSF: plasma | -0.01848 | 24 | - 0.168 | 0.131 2 | 0.9988 |
| Figure 4l | TfR$^{mu/hu}$-KI (3 months v. 10.5 months) | Two-way | % CSF: plasma | 0.0004996 | 24 | - 0.149 | 0.150 2 | 0.9999 |
| Figure 4l | 5xFAD; TfR$^{mu/hu}$-KI (1.5 months v. 3 months) | Two-way | % CSF: plasma | 0.05198 | 24 | - 0.097 | 0.201 7 | 0.8869 |
| Figure 4l | 5xFAD; TfR$^{mu/hu}$-KI (1.5 months v. 10.5 months) | Two-way | % CSF: plasma | 0.03655 | 24 | - 0.113 | 0.186 2 | 0.9724 |
| Figure 4l | 5xFAD; TfR$^{mu/hu}$-KI (3 months v. 10.5 months) | Two-way | % CSF: plasma | -0.01543 | 24 | - 0.165 | 0.134 3 | 0.9995 |
| Figure 4l | 1.5 months (TfR$^{mu/hu}$-KI v. 5xFAD; TfR$^{mu/hu}$-KI) | Two-way | % CSF: plasma | -0.04194 | 24 | - 0.191 | 0.107 7 | 0.9509 |
| Figure 4l | 3 months (TfR$^{mu/hu}$-KI v. 5xFAD; TfR$^{mu/hu}$-KI) | Two-way | % CSF: plasma | 0.02901 | 24 | - 0.120 | 0.178 7 | 0.9900 |
| Figure 4l | 10.5 months (TfR$^{mu/hu}$-KI v. 5xFAD; TfR$^{mu/hu}$-KI) | Two-way | % CSF: plasma | 0.01309 | 24 | - 0.136 | 0.162 8 | 0.9997 |
| Figure 4n | Grey Matter SMA$^{+}$ vessels (Healthy v. AD) | One-way | AU/mm$^2$ | 0.4458 | 36 | -3.068 | 3.959 | 0.9999 |
| Figure 4n | Grey Matter SMA$^{-}$ vessels (Healthy v. AD) | One-way | AU/mm$^2$ | 1.546 | 36 | -1.968 | 5.059 | 0.9201 |
| Figure 4n | Grey Matter Parenchyma (Healthy v. AD) | One-way | AU/mm$^2$ | 0.002144 | 36 | -3.511 | 3.516 | >0.9999 |
| Figure 4n | White Matter SMA$^{+}$ vessels (Healthy v. AD) | One-way | AU/mm$^2$ | 1.160 | 36 | -2.354 | 4.673 | 0.9894 |

| | | | | | | | | |
|---|---|---|---|---|---|---|---|---|
| Figure 4n | White Matter SMA$^-$ vessels (Healthy v. AD) | One-way | AU/mm$^2$ | 2.718 | 36 | -0.795 | 6.232 | 0.2665 |
| Figure 4n | White Matter Parenchyma (Healthy v. AD) | One-way | AU/mm$^2$ | 0.1815 | 36 | -3.332 | 3.695 | 0.9999 |
| Supp. Figure 3a | TfR$^{mu/hu}$-KI (1.5 months v. 3 months) | Two-way | nM | -1652 | 24 | -8762 | 5459 | 0.9777 |
| Supp. Figure 3a | TfR$^{mu/hu}$-KI (1.5 months v. 10.5 months) | Two-way | nM | -12700 | 24 | -19810 | -5589 | 1.47E-04 |
| Supp. Figure 3a | TfR$^{mu/hu}$-KI (3 months v. 10.5 months) | Two-way | nM | -11048 | 24 | -18158 | -3938 | 8.65E-04 |
| Supp. Figure 3a | 5xFAD; TfR$^{mu/hu}$-KI (1.5 months v. 3 months) | Two-way | nM | -2622 | 24 | -9732 | 4489 | 0.8597 |
| Supp. Figure 3a | 5xFAD; TfR$^{mu/hu}$-KI (1.5 months v. 10.5 months) | Two-way | nM | -11195 | 24 | -18305 | -4085 | 7.39E-04 |
| Supp. Figure 3a | 5xFAD; TfR$^{mu/hu}$-KI (3 months v. 10.5 months) | Two-way | nM | -8573 | 24 | -15684 | -1463 | 1.18E-02 |
| Supp. Figure 3a | 1.5 months (TfR$^{mu/hu}$-KI v. 5xFAD; TfR$^{mu/hu}$-KI) | Two-way | nM | -0.4553 | 24 | -7111 | 7110 | >0.9999 |
| Supp. Figure 3a | 3 months (TfR$^{mu/hu}$-KI v. 5xFAD; TfR$^{mu/hu}$-KI) | Two-way | nM | -970.5 | 24 | -8081 | 6140 | 0.9980 |
| Supp. Figure 3a | 10.5 months (TfR$^{mu/hu}$-KI v. 5xFAD; TfR$^{mu/hu}$-KI) | Two-way | nM | 1504 | 24 | -5606 | 8615 | 0.9852 |
| Supp. Figure 3b | TfR$^{mu/hu}$-KI (1.5 months v. 3 months) | Two-way | nM | -2.325 | 24 | -11.13 | 6.476 | 0.9615 |
| Supp. Figure 3b | TfR$^{mu/hu}$-KI (1.5 months v. 10.5 months) | Two-way | nM | -19.05 | 24 | -27.85 | -10.25 | 8.67E-06 |
| Supp. Figure 3b | TfR$^{mu/hu}$-KI (3 months v. 10.5 months) | Two-way | nM | -16.73 | 24 | -25.53 | -7.928 | 6.16E-05 |
| Supp. Figure 3b | 5xFAD; TfR$^{mu/hu}$-KI (1.5 months v. 3 months) | Two-way | nM | -3.094 | 24 | -11.89 | 5.707 | 0.8818 |
| Supp. Figure 3b | 5xFAD; TfR$^{mu/hu}$-KI (1.5 months v. 10.5 months) | Two-way | nM | -15.38 | 24 | -24.18 | -6.579 | 1.97E-04 |
| Supp. Figure 3b | 5xFAD; TfR$^{mu/hu}$-KI (3 months v. 10.5 months) | Two-way | nM | -12.29 | 24 | -21.09 | -3.486 | 2.87E-03 |
| Supp. Figure 3b | 1.5 months (TfR$^{mu/hu}$-KI v. 5xFAD; TfR$^{mu/hu}$-KI) | Two-way | nM | -0.6930 | 24 | -9.494 | 8.108 | 0.9998 |
| Supp. Figure 3b | 3 months (TfR$^{mu/hu}$-KI v. 5xFAD; TfR$^{mu/hu}$-KI) | Two-way | nM | -1.462 | 24 | -10.26 | 7.339 | 0.9951 |
| Supp. Figure 3b | 10.5 months (TfR$^{mu/hu}$-KI v. 5xFAD; TfR$^{mu/hu}$-KI) | Two-way | nM | 2.980 | 24 | -5.821 | 11.78 | 0.8970 |
| Supp. Figure 3c | TfR$^{mu/hu}$-KI (1.5 months v. 3 months) | Two-way | nM | -2.161 | 24 | -13.44 | 9.118 | 0.9905 |
| Supp. Figure 3c | TfR$^{mu/hu}$-KI (1.5 months v. 10.5 months) | Two-way | nM | -23.37 | 24 | -34.65 | -12.10 | 1.71E-05 |
| Supp. Figure 3c | TfR$^{mu/hu}$-KI (3 months v. 10.5 months) | Two-way | nM | -21.21 | 24 | -32.49 | -9.935 | 7.16E-05 |
| Supp. Figure 3c | 5xFAD; TfR$^{mu/hu}$-KI (1.5 months v. 3 months) | Two-way | nM | -3.336 | 24 | -14.61 | 7.942 | 0.9388 |
| Supp. Figure 3c | 5xFAD; TfR$^{mu/hu}$-KI (1.5 months v. 10.5 months) | Two-way | nM | -15.78 | 24 | -27.06 | -4.504 | 2.80E-03 |
| Supp. Figure 3c | 5xFAD; TfR$^{mu/hu}$-KI (3 months v. 10.5 months) | Two-way | nM | -12.45 | 24 | -23.72 | -1.167 | 2.46E-02 |
| Supp. Figure 3c | 1.5 months (TfR$^{mu/hu}$-KI v. 5xFAD; TfR$^{mu/hu}$-KI) | Two-way | nM | -0.2429 | 24 | -11.52 | 11.04 | 0.9999 |
| Supp. Figure 3c | 3 months (TfR$^{mu/hu}$-KI v. 5xFAD; TfR$^{mu/hu}$-KI) | Two-way | nM | -1.419 | 24 | -12.70 | 9.860 | 0.9986 |
| Supp. Figure 3c | 10.5 months (TfR$^{mu/hu}$-KI v. 5xFAD; TfR$^{mu/hu}$-KI) | Two-way | nM | 7.349 | 24 | -3.929 | 18.63 | 0.3635 |

**Table S5.** Demographics of human AD patients and age-matched healthy controls.

| **Sample ID** | **Sex** | **Age** | **Diagnosis** |
|---|---|---|---|
| 531677A(3) | F | 84 | normal tissue |
| 531653A(*12*) | M | 81 | normal tissue |
| 531621A(*12*) | F | 84 | normal tissue |
| 531060A(2) | M | 84 | normal tissue |
| 532036A(3) | F | 96 | AD |
| 531885A(3) | F | 87 | AD |
| 531231A(9) | F | 90 | AD |
| 531200A(2) | F | 80 | AD |
